# Age-associated changes in endothelial transcriptome and chromatin landscape correlate with elevated risk of hemorrhage

**DOI:** 10.1101/2023.02.10.528012

**Authors:** Kshitij Mohan, Gilles Gasparoni, Abdulrahman Salhab, Michael M. Orlich, Robert Geffers, Steve Hoffmann, Ralf H. Adams, Jörn Walter, Alfred Nordheim

## Abstract

Intracerebral hemorrhage (ICH), a devastating form of stroke, is a leading global cause of human death and disability. The major risk factors for ICH include increasing age, hypertension and cerebral amyloid angiopathy. Despite high mortality and morbidity associated with ICH, the mechanisms leading to blood-brain barrier (BBB) dysfunction with age and development of hemorrhagic stroke is poorly understood. In the vasculature of the central nervous system, endothelial cells (ECs) constitute the core component of the BBB and provide a physical barrier due to tight junctions, adherens junctions, and basement membrane layers. In this study, we show in brains of mice that incidents of intracerebral bleeding increase with advancing age. After isolation of an enriched population of cerebral ECs, we studied gene expression in ECs isolated from murine brains of increasing ages of 2, 6, 12, 18, and 24 months. The study reveals age-dependent dysregulation of 1388 genes in the ECs, including many involved in the maintenance of BBB and vascular integrity. Since epigenetic mechanisms regulate gene expression, we also investigated age-dependent changes at the levels of CpG methylation and accessible chromatin in cerebral ECs. Our study reveals correlations between age-dependent changes in chromatin structure and gene expression. We find significant age-dependent downregulation of the apelin receptor (*Aplnr*) gene along with an age-dependent reduction in chromatin accessibility of the promoter of this gene. *Aplnr* is known to play a crucial role in positive regulation of vasodilation and is implicated in vascular health. Interestingly, we also observe an age-dependent reduction in the protein expression levels of the apelin receptor in the brain, potentially implicating the apelin receptor to be critical for the increased risk of intracerebral hemorrhage with ageing.

## Introduction

Intracerebral hemorrhage (ICH) is a leading cause of human death and disability, with 3.4 million new incidents and 3.2 million deaths reported globally every year^1^. Among surviving patients, a majority suffers from disabilities, with only less than 40% survivors attaining full functional recovery^2^. Age is the most significant risk factor for ICH, reaching a maximum in individuals older than 75 years. ICH incidences in the age groups of 35-54 years, 55-74 years and 75-94 years have been reported to amount to 5.9, 37.2, and 176.3 per 100,000 individuals, respectively^3^. ICH is caused by the rupture of small to medium-sized blood vessels with a diameter of 100 – 600 µm and is typically a manifestation of cerebral small vessel disease^4^. Ageing leads to structural, functional, and mechanical changes in small blood vessels, resembling the changes in small vessels arising from chronic hypertension. These changes lead to the degeneration of the vascular wall, causing development of small aneurysms and microbleeds in the deeper structures, indicative of risk for intracerebral hemorrhages^5, 6^.

The cerebral microvasculature has the unique property of a highly selective blood-brain barrier (BBB) playing an essential role in brain homeostasis by tightly regulating paracellular and transcellular transport of ions, macromolecules, pathogens, and cells of the immune system between blood and brain^7^. The central nervous system’s microvasculature comprises a continuous monolayer of ECs forming the blood vessels’ inner-most layer, pericytes that wrap around the ECs, basement membrane (BM) surrounding the vascular tube, astrocytes, and neurons^8^. The endothelial monolayer forming the innermost wall of the cerebral microvasculature constitutes the core component of the BBB. The presence of continuous tight junctions (TJs) that form a physical barrier between the ECs, the adherens junctions (AJs), and the basement membrane (BM) are the major structures that impart on the cerebral microvasculature its unique characteristic of BBB function^9, 10^. Cytoskeletal activities and cell-cell interactions of ECs are regulated to an important extent by the transcription factor SRF^11-15^. In this study, however, we did not impair SRF activity experimentally, but rather asked whether the aging process impinges on BBB function, potentially via modulating endogenous SRF activity.

Increasing age is associated with endothelial dysfunction, as well functional, structural, and mechanical changes in the blood vessels. While age is a non-modifiable risk factor for several vascular diseases, including intracerebral hemorrhage^6, 16^, age-dependent changes in expression in cerebral ECs are still insufficiently characterized. In this study, we have used RNA-seq to characterize the age-dependent changes in mRNA expression levels in ECs isolated from murine brains of increasing ages of 2, 6, 12, 18, and 24 months.

Further, we interrogated the possibility of age-associated epigenetic regulation of cerebral EC gene expression, primarily focusing on methylation of cytosines in CpG-rich promoter regions of genes^17^. We performed reduced representation bisulfite sequencing (RRBS), a bisulfite conversion-based protocol that enriches CG-rich regions of the genome, to study age-dependent changes in CpG methylation at promoter regions in the cerebral ECs isolated from murine brains of increasing age. We also performed transposase-accessible chromatin sequencing (ATAC-seq). Our study aims to understand age-dependent cerebral bleeding better and expand our knowledge of intracerebral hemorrhage, with potential relevance to human patients.

## Methods

We studied the effect of ageing on the incidence of spontaneous cerebral bleeding in wild-type mice and studied the underlying age-associated changes in (a) gene expression using RNA-seq, (b) genomic DNA methylation levels using Reduced Representation Bisulfite Sequencing (RRBS) and (c) chromatin landscape using Assay for Transposase-accessible Chromatin Sequencing (ATAC-seq) in cerebral Endothelial Cells (ECs). A detailed description of all the methods is included in the Supplementary Information.

## Results

### Cerebral bleedings increase with age in mice

Brains were dissected from C57Bl6 wild-type mice and C57Bl6 mice with floxed, yet non-recombined alleles of the *Srf* and *Mrtf* loci. These floxed mice do not show any difference in phenotype compared to wild-type mice, and have been used as controls, referred to as ‘control mice’ henceforth. We find that both wild-type mice and control mice (C57Bl6 mice with floxed, yet non-recombined alleles of the *Srf* and *Mrtf* loci) of different ages show an age-dependent increase in bleedings. To quantify the cerebral bleedings, we adhered to the bleedings that had diameter between 50 and 300 µm based on available literature^18, 19^. Notably, the bleedings were of variable sizes (diameter between 50 and 250 µm) and not restricted to any specific brain region **(Figure 1B)**. To quantify the number of bleedings in each brain, we evaluated every tenth slide of hematoxylin eosin(H&E)-stained brain sections, collectively yielding 20 slides per brain for each animal.

**Figure 1.**
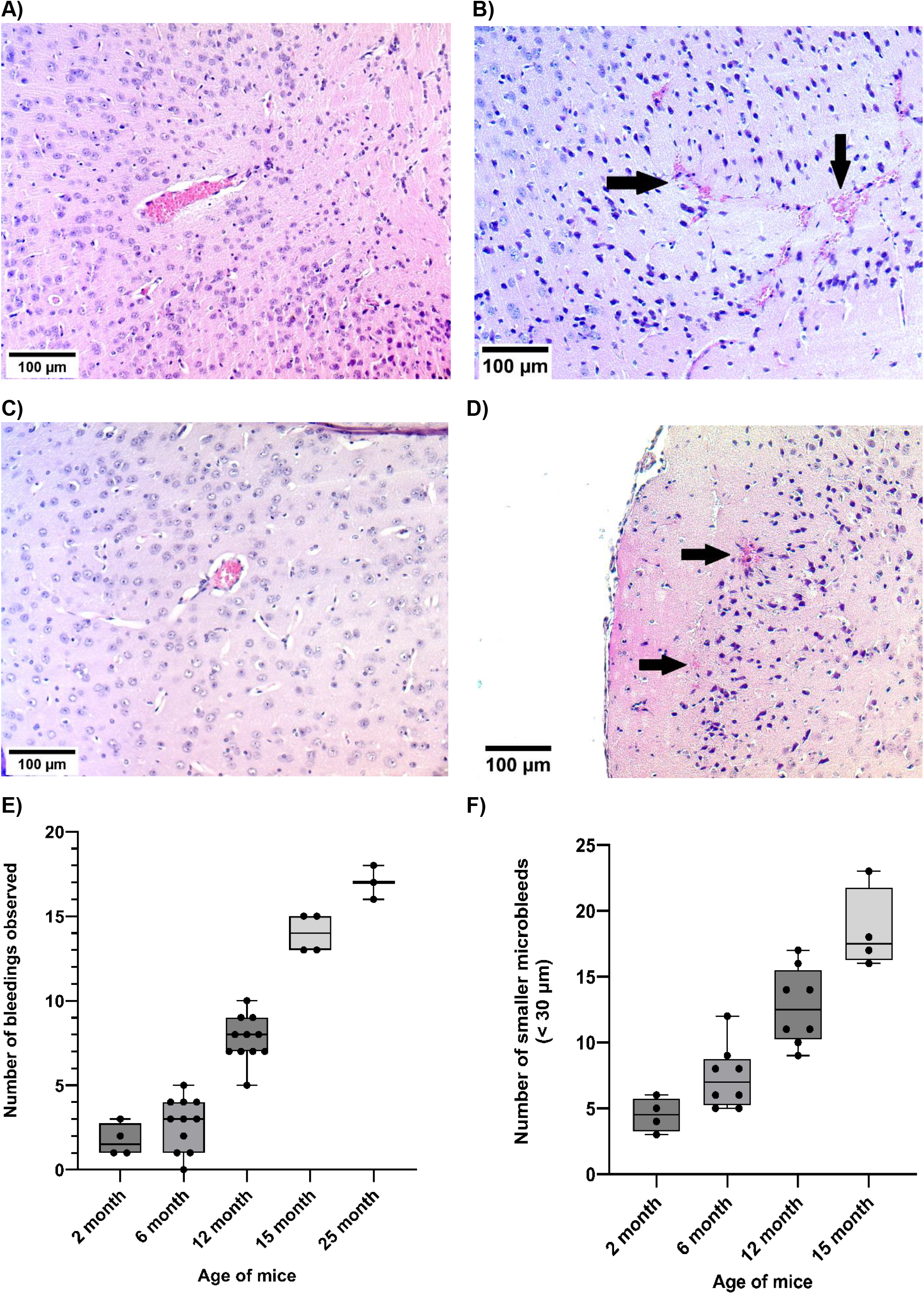
The number of bleedings in the brain of mice increases with age. **(A)** An intact blood vessel. **(B)** Cerebral bleedings from a blood vessel and the leakage of erythrocytes (arrows) stained pink by H&E stain in the brain parenchyma. **(C)** An intact smaller blood vessel in the mouse brain. **(D)** Smaller microbleeds from a blood vessel and the leakage of erythrocytes (arrows) stained pink by H&E stain in the brain parenchyma. **(E)** Quantification of bleedings indicates an age-dependent increase in the number of bleedings in the brain. The average number of bleedings recorded per brain in 2 months-old mice (number of mice, n = 4) is 1.75. In 6 months (n= 11), 12 months (n=11), 15 months (n=4) and 25 months-old mice (n=3), the average numbers of bleeding recorded are 2.73, 7.72, 13.75 and 17, respectively. Error bars represent mean ± se (standard error of the mean). **(F)** Quantification indicates an age-dependent increase in the number of microbleeds in brain. The average number of microbleeds recorded (every 10th slide was quantified) per brain in 2 months-old mice (number of mice, n = 4) is 4.5. In 6 months (n= 8), 12 months (n=8) and 15 months-old mice (n=4), the average numbers of microbleeds (every 10th slide was quantified) are 8.75, 13 and 18.5, respectively. Error bars represent mean ± se (standard error of the mean).

**Figure 2.**
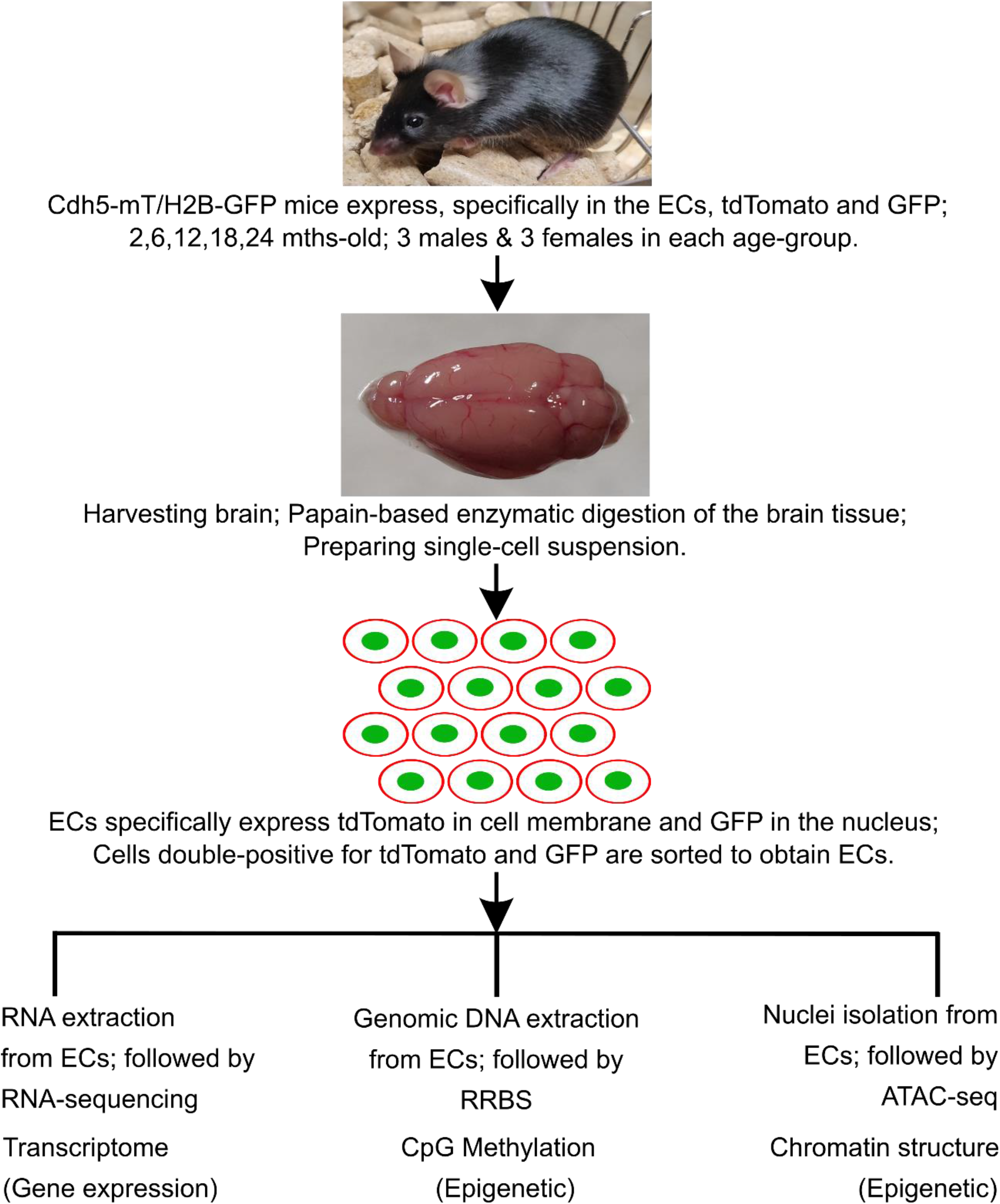
Experimental design to study age-associated transcriptomic and epigenetic changes in the cerebral ECs. Cdh5-mT/H2B-GFP transgenic mice that specifically express – in the ECs – membrane targeted tandem dimer Tomato fluorescence and H2B-GFP, thereby expressing Tomato fluorescence in cell membrane and GFP in the nucleus, were used to isolate pure population of cerebral ECs. 3 male and 3 female mice from each of the 2,6,12,18 and 24 months-old cohort were used to for the study. RNA-seq, RRBS and ATAC-seq were performed to study the age-associated transcriptomic and epigenetic changes (CpG methylation and chromatin accessibility) in the cerebral ECs.

The average number of bleedings recorded per brain in two-month-old mice (number of mice used in this age-group, n = 4) was 1.75 with a standard deviation (sd) of 0.96. In 6 month (n= 11), 12 month (n=11), 15 month (n=4) and 25-month-old mice (n=3), the average numbers of bleedings increased to 2.73 (sd = 1.56), 7.72 (sd = 1.35), 13.75 (sd = 0.96) and 17 (sd = 1), respectively **(Figure 1E)**. While there was no significant increase in the number of bleedings in the brains of the 6-month-old mice compared to the 2-month-old mice (adjusted p-value = 0.73), we noted a significant increase in the bleedings occurring at 12 months and 15 months compared to 2-month-old brains (adjusted p-value < 0.001). However, we did not observe a significant increase in the incidents of cerebral bleedings between 15 and 25 months (ANOVA followed by Tukey’s HSD). Since H&E staining used to quantify bleedings in our study specifically detect fresh cerebral bleedings^20^, we argue that the increase in the number of bleedings observed with advancing age in mice is not merely accumulation of old bleedings over time but rather indicates fresh reflects an increase of the incidents of new bleedings with advancing age.

### Small-size cerebral microbleeds increase with age in mice

After observing an age-dependent increase in the average number of bleedings per brain in mice, we also observed smaller microbleeds and performed a second study focusing on these smaller microbleeds, characterized by smaller size of bleedings (diameter < 30 µm). The brains dissected from C57Bl6 wild-type and control mice of different ages also revealed an increased frequency of smaller microbleeds with ageing. The number of unique bleedings in a mouse brain was quantified by recording the blood vessels with extravasated erythrocytes that were stained pink in the H&E staining or by observing the leakage of erythrocytes into the brain parenchyma **(Figure 1D)**. To quantify the number of small-sized microbleeds in each brain in this study, we evaluated every tenth slide, yielding 20 slides per brain for each animal. The average number of microbleeds recorded per brain, quantified in serial sections using every 10^th^ slide, in the 2-month-old mice (number of mice used in this age-group, n = 4) was 4.5 with a standard deviation (sd) of 1.29. With six months (n= 8), 12 months (n=8) and 15 months (n=4), the average numbers of bleeding recorded were8.75 (sd = 2.39), 13 (sd = 2.62) and 18.5 (sd = 3.11), respectively **(Figure 1F)**. While there was no significant increase in the number of bleedings in the brains at 6 months mice compared to 2 months (adjusted p-value = 0.29, ANOVA followed by Tukey’s HSD), we found a significant increase in the bleedings that occur in 12- and 15-month-old mice compared to the 2-month-old mice (adjusted p-value < 0.001). Since brain sections from 25-month-old samples were not available for analysis of the smaller cerebral microbleeds, we only quantified the number of smaller microbleeds in 2-, 6-, 12- and 15-month-old age groups. Also, we did not quantify the number of microbleeds in 3 samples belonging to 12 and 15 months-old groups, accounting for the difference in the total number of brains examined for bleedings and smaller microbleeds.

### Purification of cerebral ECs from Cdh5-mT/H2B-GFP mice

To purify endothelial cells from the brain vasculature, we took advantage of transgenic *Cdh5-mT/H2B-GFP* mice. Driven by the pan-endothelial vascular endothelial cadherin (Cdh5) promoter, these mice express membrane-targeted tdTomato (mT) fluorescent protein in cell membranes and green fluorescent protein fused to histone2B (H2B-GFP) in nuclei. To characterize age-associated transcriptomic and epigenetic changes in the cerebral ECs, three male and three female mice were used belonging to each of the 2-, 6-, 12-, 18- and 24-month age groups. We also measured the body weight and weights of different organs but did not observe any age-associated changes **(Figure S1, Table S1)**. When we analyzed single-cell suspensions prepared from transgenic animals using FACS, approximately 7% of the cells were positive for both GFP and tdTomato. A double-positive GFP^+^/tdTomato^+^ cell population was absent in the single-cell suspension prepared from wild-type, non-transgenic mouse brains. To ensure the highest purity of ECs in the isolated population, we used FACS to sort cells. The GFP^+^/tdTomato^+^ double-positive cells were sorted again for GFP^+^/tdTomato^+^, a process known as reanalysis, where we determined the yield of ECs in the sorted cells to be approximately 99%. **(Figure 3A,3C, Figure S2)**.

**Figure 3.**
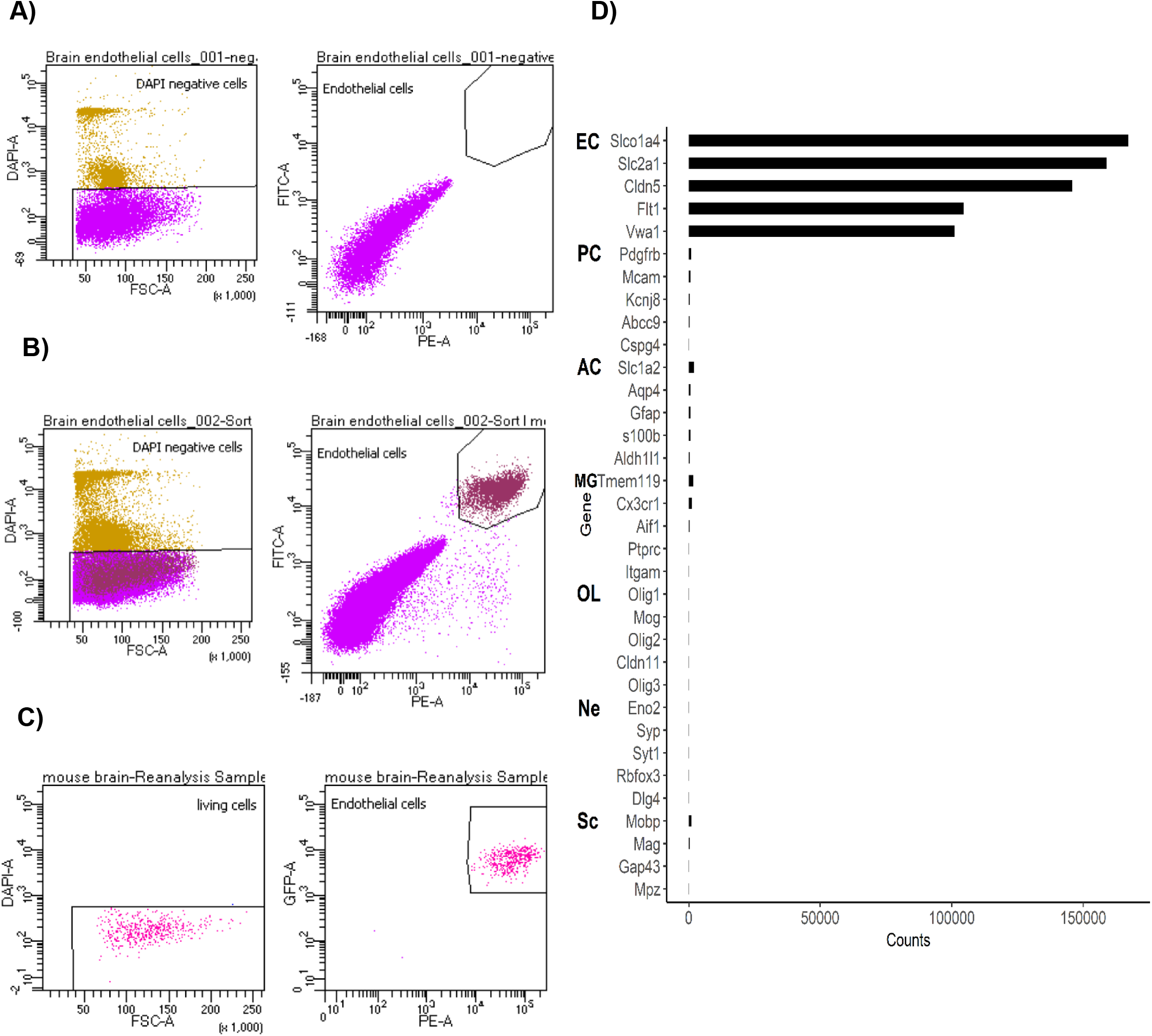
FACS profile of sorted ECs. **(A)** The single-cell suspension prepared from a wild-type mouse brain does not show a population of cells positive for GFP and tdTomato, **(B)** while the singlecell suspension prepared from Cdh5-mT H2B-GFP transgenic mouse brain shows the presence of EC population double positive for GFP and tdTomato fluorescence. **(C)** The double positive cells (GFP+ tdTomato+) were sorted. The purity of sorted ECs was ~ 99% as confirmed by the reanalysis of the sorted cells. **(D)The relative transcript levels of specific markers for various cells of the neurovascular unit and the brain**. The specific markers for endothelial cells (ECs) are very highly enriched in comparison to the markers for other cells of the neurovascular unit, confirming the purity of the ECs sorted. (EC – endothelial cells, PC – pericytes, AC – astrocytes, MG – microglia, OL – oligodendrocytes, Ne – neurons, Sc – Schwann cells)

The purity of the isolated endothelial cell population was further confirmed using data obtained from RNA-sequencing of sorted GFP^+^/tdTomato^+^ cells by comparing the relative transcript levels of specific markers for endothelial cells (*Slco1a4, Slc1a2, Cldn5, Flt1*, and *Vwa1*), pericytes (*Pdgfrb, Mcam, Abcc9, Kcnj8*, and *Cspg4*), astrocytes (*Slc1a2, Aqp4, Gfap, S100b*, and *Aldh1l1*), microglia (*Tmem119, Cx3cr1, Aif1, Ptprc*, and *Itgam*), oligodendrocytes (*Olig1, Mog, Olig2, Cldn11*, and *Olig3*), neurons (*Eno2, Syp, Syt1, Rbfox3*, and *Dlg4*) and Schwann cells (*Mobp, Mag, Gap43*, and *Mpz*). To choose specific markers for different cell types in brain, we used the molecular atlas of cell types in the brain vasculature^21^. As expected, transcripts of EC-specific markers were very highly expressed (normalized counts> 100,000) in sorted cells. In contrast, the expression of transcripts of markers of other cells of the neurovascular unit and neural cells were low (normalized counts ~ 1000) to absent, confirming the high level of purity of our EC population. **(Figure 3D)**

### RNA-seq reveals age-dependent dysregulation of gene expression in cerebral ECs

To study age-dependent transcriptomic changes in cerebral ECs, we performed RNA-seq of ECs purified from brains of three males and three females belonging to each of the 2-, 6-, 12-, 18- and 24-month-old age groups. We performed linear regression analysis on RNA-seq data for the thirty samples across all time points (2, 6, 12, 18 and 24 months), adjusting for the sex-specific effects, to identify genes dysregulated in ageing. After adjusting for multiple testing using the false discovery rate (FDR; Benjamini-Hochberg), 1388 genes were found to be dysregulated with increasing age. Of these, 675 genes were downregulated while 713 were upregulated with age **(Excel S1)**. The heatmap of significantly dysregulated genes reveals distinctive, age-group-specific expression patterns **(Figure 4A, Figures S3**,**S4)**. Enrichment analysis performed on the top 1000 genes dysregulated with age in the cerebral ECs reveal 345 GO processes that are enriched **(Excel S2)**. The processes ‘cardiovascular system development’ (GO:0072358), ‘vasculature development’ (GO:0001944), ‘regulation of blood vessel endothelial cell migration’ (GO:0043535), ‘blood vessel morphogenesis’ (GO:0048514) and ‘regulation of endothelial cell migration’ (GO: 0010594) – associated with cerebral vasculature – were among the enriched processes. We also found the enrichment of processes associated with actin dynamics, cytoskeleton organization, and Cdc42 signaling pathway. We also performed enrichment analysis on the top 100 and top 500 genes and observed enrichment in 67 and 158 GO processes, respectively. **(Excel S3, S4, Figures S5, S6)**.

**Figure 4.**
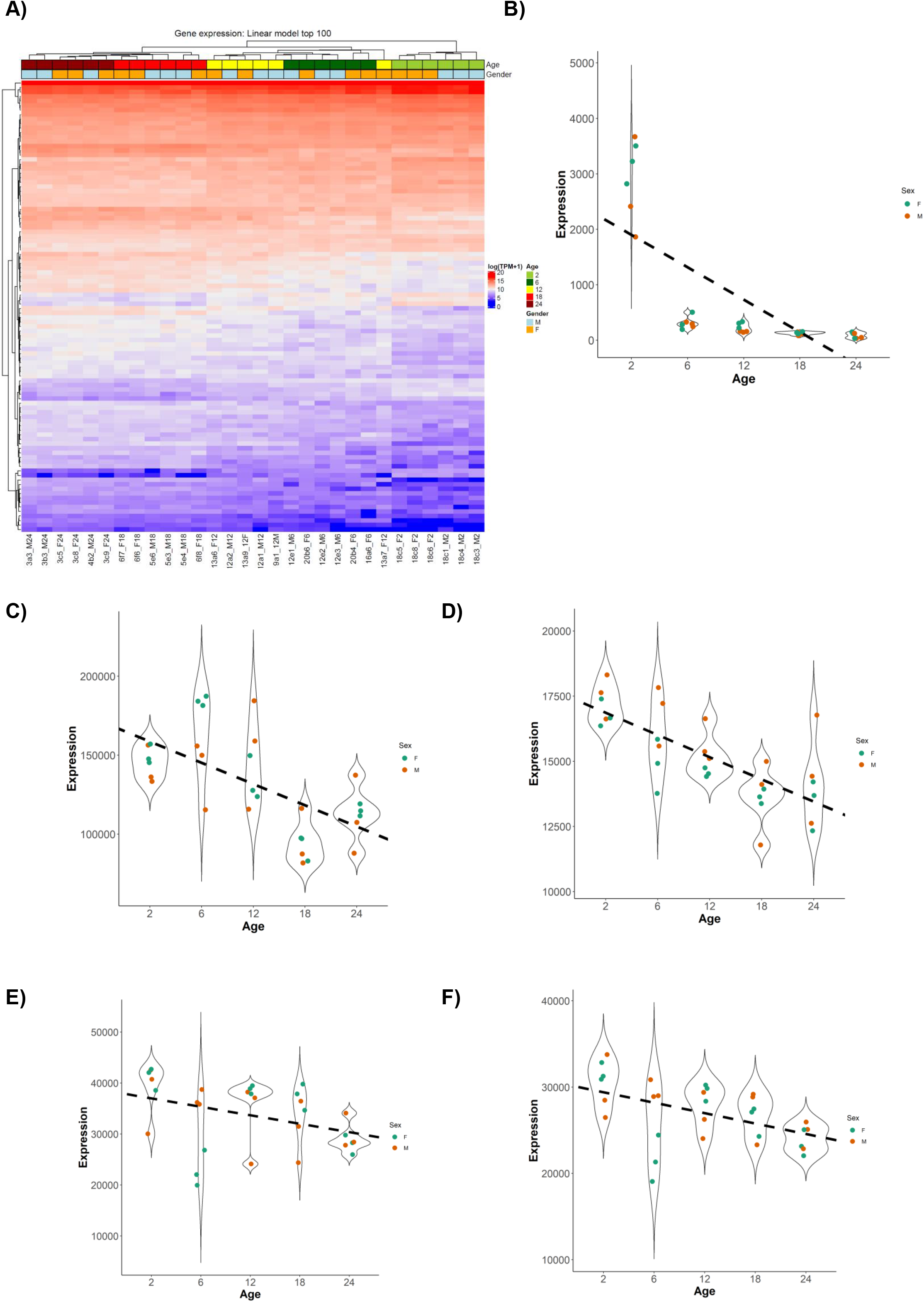
(A) Heatmap clustering based on top 100 genes that exhibit an age-dependent dysregulation in the linear regression model. **(B) The expression level of *Aplnr* with age**. Linear regression analysis suggests that the expression of *Aplnr* decreases with age, with a correlation coefficient (r-squared value) of 0.47 (adjusted p-value = 0.004). The plot showing the normalized counts of *Aplnr* in each sample indicates the downregulation of *Aplnr* is the sharpest between the ages of 2 months and 6 months. TPM : Transcripts per million. **Expression of genes encoding key components of tight junctions across age**. The violin plots with linear regression analysis indicates that among major tight junction proteins at the BBB, **(C)** *Cldn5* and **(D)** *F11r* show an age-dependent downregulation, with correlation coefficients of 0.42 (adjusted p-value = 0.009) and 0.59 (adjusted p-value = 0.0008), respectively. However, no age-dependent downregulation of other tight junction components such as **(E)** *Ocln* and **(F)** *Jam3* was observed (adjusted p-values > 0.1).

In our RNA-seq dataset, we focused on the expression patterns of genes involved in the GO processes associated with vasculature development and cytoskeletal dynamics. We then searched specifically for expression patterns of (i) genes implicated in hypertension and associated with human cerebral small vessel diseases, (ii) genes encoding structural components associated with the ECs in blood-brain barrier, and (iii) genes involved in the maintenance of vascular integrity in the brain. We identified both the *Aplnr* gene, encoding Apelin receptor, and the gene encoding its ligand Apelin (*Apln*) to be strongly downregulated with age. In zebrafish, the Apelin signaling pathway plays a major role in angiogenesis by regulating angiogenic sprouting, tip-cell morphology, and the EC sprouting behavior. Interestingly, fish embryos lacking either the apelin ligand or the apelin receptor have been shown to suffer from impaired inter-segmental vessel formation^22^. *Aplnr* is also known to play a crucial role in positive regulation of vasodilation; and an increased hypertension in mice, rats and humans have been associated with reduced serum Apelin and disruption of the Apelinergic axis^23-25^. *In vitro* studies have established the role of Apelin receptor in regulating biomechanical and morphological properties of ECs by regulating signaling pathways that mediate adaptation of ECs to the flow conditions by modulating EC morphology, elasticity, adhesion, and spreading^26^. We observed that both *Aplnr* and *Apln* display a significant, progressive decrease in expression with age in cerebral ECs, as indicated by pair-wise comparison of different age groups. The log2-fold change values of *Aplnr* were – 3.18, – 4.10, – 4.51, and – 5.24 (adjusted p-values < 0.001) in 6-, 12-, 18-, and 24-month-old male mice, respectively – as compared to 2-month-old mice. In females, the log2-fold change values of *Aplnr* in the females of 6, 12, 18, and 24 months were – 3.30, – 3.47, – 4.54, and – 4.98 (adjusted p-values < 0.001), respectively. Thus, *Aplnr* transcript levels in the cerebral endothelial cells continue to steadily decline during aging from a ~10-fold change (2 vs 6) to a ∼32-fold change (2 vs 24) even after six months of age **(Figure 4B)**. The Apelin signaling pathway is downstream of Notch signaling, with Notch signaling negatively regulating Apln^22^. In zebrafish and HUVECs, inhibition of Notch resulted in the upregulation of Apln while activation of Notch led to the inhibition of Apln^22^. We then focused on the Apelin signaling pathway (KEGG:mmu04371) and compared the expression of relevant pathway genes to our RNA-seq data. Interestingly, we observed that *Notch3* was significantly upregulated with age.

Next, we attempted to identify age-dependent changes in transcripts of genes encoding structural components in the BBB endothelium. Among the tight junction components, we found downregulation of the *Cldn5* gene, the major claudin expressed in the ECs of the CNS. The linear regression analysis suggests age-dependent downregulation of the *Cldn5* gene (adjusted p-value = 0.009) with a correlation coefficient (r-squared value) of 0.42, meaning that 42% of the changes in the expression level of Cldn5 gene can be correlated with ageing. We did not observe any significant age-dependent dysregulation of other claudin isoforms, i.e., *Cldn1, Cldn2, Cldn3, Cldn10, Cldn11*, and *Cldn12*. Also, no age-dependent dysregulation was observed for *Ocln* (occludin), another key component of the tight junctions at BBB. Among the junctional adhesion molecules, we found the *F11r* gene encoding the JAM-A protein to be significantly downregulated. Other critical junctional adhesion molecules implicated to be vital for the BBB, such as *Jam2, Jam3*, and *Igsf5*, however, were not significantly dysregulated. **(Figure 4C)**

Furthermore, we observed a significant age-dependent downregulation of *the Cdh2* gene (N-cadherin) in cerebral ECs but no age-dependent dysregulation of *Cdh5* (VE-cadherin), the major component of adherens junctions (AJ) in the cerebral ECs. Also, no significant age-dependent change in the expressions of other AJ components such as *Cdh1*, β-Catenin (*Ctnnb1*), α-Catenin (*Ctnna1*), p120 (*Ctnnd1*), and plakoglobin (*Jup*) was observed **(Figure S7)**. The enrichment analysis based on top 1000 genes dysregulated with age in the ECs suggests an enrichment in the processes associated with ‘cytoskeleton organisation’ (GO: 0007010), ‘actin cytoskeleton organisation’ (GO:0030036), ‘negative regulation of cytoskeleton organization’ (GO:0051494), ‘actin filament-based process’ (GO:0030029), ‘Cdc42 protein signal transduction’ (GO:0032488). Among the genes associated with actin cytoskeleton, we observe a significant age-associated downregulation of *Cdc42* (adjusted p-value = 0.005) and *Cdc42se1* (adjusted p-value = 0.003). We also observed a significant downregulation in the expression of *Arf1* and *Arf6* genes involved in maintaining the actin cytoskeleton dynamics by acting downstream to Cdc42 in the actin polymerization pathway. However, no age-dependent dysregulation was observed in *RhoA, Rac1, Vcl, Vasp, Anln*, and *Actn4*, encoding various other actin-binding proteins. Among the key cytoplasmic components of the tight junction complex, *Tjp1, Tjp2, Tjp3, Cgn*, and *Afdn* were not dysregulated with age **(Figure S8)**. We did not observe any significant age-dependent dysregulation of major scaffolding proteins present at the BBB such as *Magi3, Magi1, Patj, Mpp1, Mpp5, Mpp7, Dlg1, Mpdz, Pard3, Pard6a*, and *Pard6b* **(Figure S9)**. Regarding genes that encode basement membrane and extracellular matrix components responsible, we did not observe any significant age-dependent dysregulation of *Col4a1* and *Col4a2*. Among the laminins, no significant age-dependent dysregulation of *Lama2, Lama5, Lamb1*, and *Lamc1*, the major laminins present at the extracellular matrix, was observed. Also, there were no significant age-dependent changes in the gene expression levels of *Nid1* (Nidogen1), *Nid2* (Nidogen2), or isoforms of integrins α*v*β*3*, α*5*β*1*, α*6*β*1*, α*1*β*1*, α*6*β*4 and* α*v*. **(Figure S10)**

In human patients, a loss-of-function mutation in the *HTRA1* gene leads to the development of CARASIL, a rare cerebral small vessel disease, in which patients display stroke-like symptoms. The *HTRA1* gene encodes a serine protease that regulates TGF-β signaling necessary to maintain vascular integrity in the brain. We observed a significant age-dependent downregulation of *Htra1* in cerebral murine ECs, as revealed by the linear regression analysis of our RNA-seq data. Pairwise-comparison suggests a significant downregulation of the *Htra1* gene in the 18 and 24 months-old males compared to the 2 months-old group. The log2-fold change of *Htra1* in 18 months-old mice was – 1.5 (adjusted p-value < 0.001) and – 1.31 in 24 months-old mice (adjusted p-value < 0.001), indicating approximately 3-fold downregulation in the expression of *Htra1* in the cerebral ECs of old mice. We also observed a significant age-dependent downregulation of the *Mfsd2a* gene, encoding major facilitator super family domain containing 2a protein critical for the formation and maintenance of the BBB^27^.

### DNA Methylation screen in cerebral ECs

Next, we examined potential age-associated epigenetic changes in murine ECs using reduced representative bisulfite sequencing (RRBS). We observed no major global methylation changes upon ageing except for a small increase in the oldest mice **(Figure 5A, Figure S9)**. To identify local age-dependent methylation changes, we applied a 1kb tiling approach to aggregate single CpG methylation values and used a linear regression model on all obtained 164,020 tiles. For the top 1000 ranking tiles according to p-value (age) we found more tiles (710/1,000) with increased methylation in old mice (hypermethylation), and only 210/1,000 showed decreased methylation levels (hypomethylation). As a sanity check, we clustered samples using the top 1,000 age-associated CpGs and obtained an apparent clustering according to age **(Figure 5B)**. Across the 710 top hypermethylated tiles the median methylation difference between youngest mice (2 months) and oldest mice (18 months) was 9.8% **(Figure 5C)** with a minimum difference of 0.5% and a maximum of 76.7% while we observed similar metrics for the 210 hypomethylated tiles (median: 13.8%, min: 0.15%, max: 69.9%).

**Figure 5.**
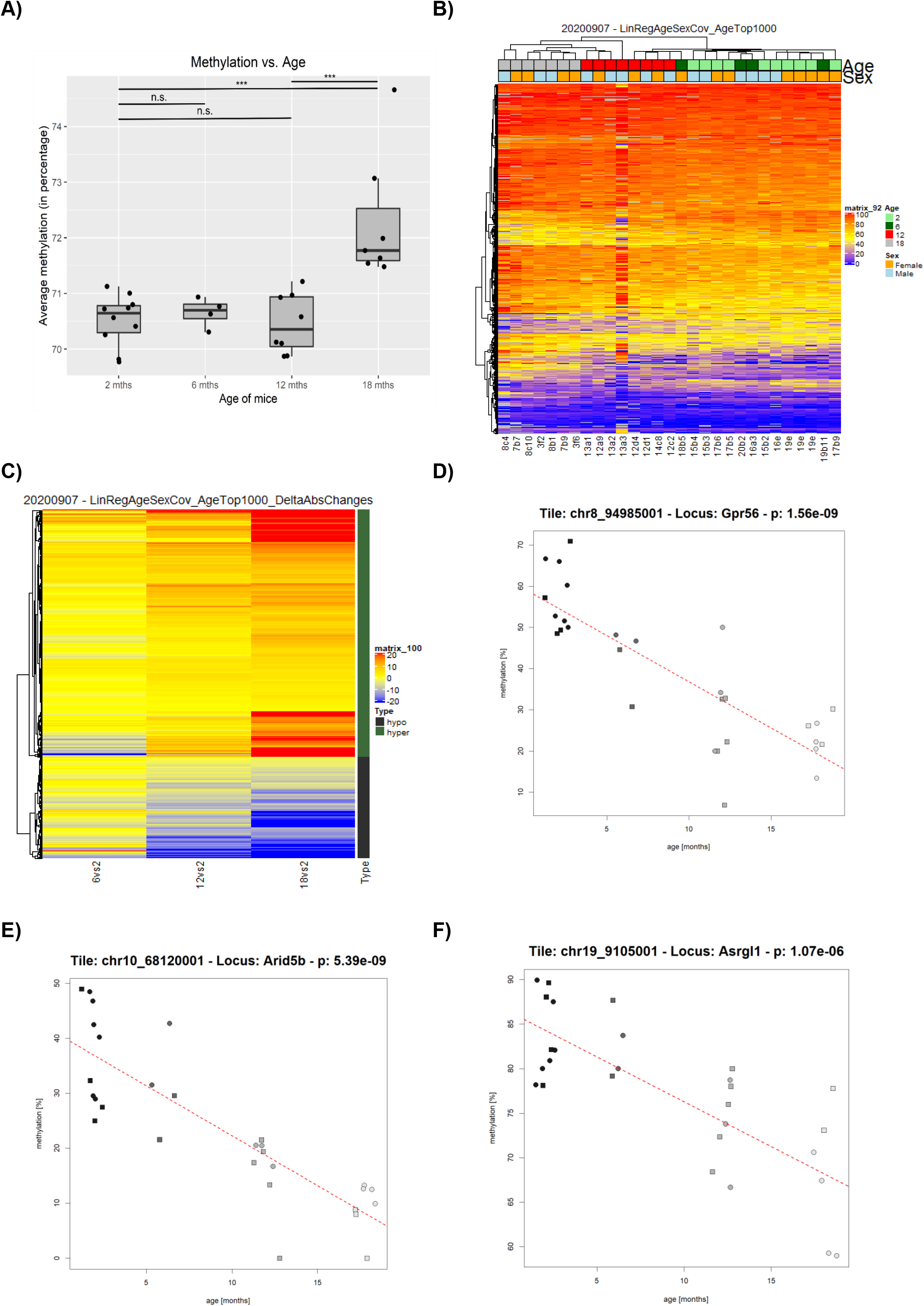
(A) ECs isolated from the brains of 18 months-old mice show increased methylation level compared to the 2, 6 and 12 months-old mice. The average methylation level across all the CpGs investigated by RRBS was 70.52 % (sd = 0.46) in the 2 months-old mice, 70.66% (sd = 0.27) in 6 months-old, 70.46% (sd = 0.53) in 12 months-old and 72.30% (sd = 1.17) in 18 months-old mice. The average methylation level of CpGs in the ECs isolated from 18 months-old mice is significantly higher in comparison to the 2, 6 and 12 months-old mice (adjusted p-values < 0.01). However, no significant change in the level of methylation is observed in the 6 or 12 months-old mice in comparison to the 2 months-old mice. Error bars represent mean ± se (standard error of the mean). One-way analysis of variance (ANOVA), followed by Tukey’s HSD test were performed to compare all the possible pairs. (**n.s**. : non-significant, **\*\*\*** : p < 0.001) **(B)** Heatmap clustering based on top 1,000 CpGs associated with ageing in the linear regression model. **(C)** Absolute methylation differences for 2 months-old versus 6, 12 and 18 months-old cohorts, respectively. To enhance visualization color gradients were capped at -20% and 20%. **(D-F)** Methylation dynamics across age groups for the top 3 ranking tiles from the linear regression analysis. The top 3 ranking tile with the most significant differential methylation dynamics with age were associated with **D)** *Gpr56*, **E)** *Arid5b* and **F)** *Asrgl1* genes. To avoid overplotting data points were subjected to a slight jitter on the x-axis. Tile location, related gene locus and obtained p value from the linear model are indicated in the respective header.

As a result, we identified a total of 12 significant tiles after applying FDR correction **(Excel S5)**. Ten out of these twelve were associated with age-associated hypomethylation, while only two showed hypermethylation. Annotation of the top significant tile **(Figure 5D)** to its closest gene revealed *Gpr56*, a gene encoding for a G protein-coupled receptor previously linked to brain development^28^ and multiple sclerosis^29^. The second top-ranking tile was linked to the *Arid5b* gene for which ageing effects previously were shown in monocytes^30^. Besides, methylation dysregulation for *Arid5b* was reported in a large epigenetic screen for epilepsy^31^ but also in human postmortem brain tissue from Alzheimer’s Disease patients^32^. Also, several of the other tiles could be linked to genes previously linked to neurological phenotypes, ageing, or age-related diseases. Among them are *Camta1*, which has been implicated in Alzheimer’s Disease^33^, age-related macular degeneration^34^, and Purkinje cell degeneration^35^; *Degs2*, implicated in schizophrenia^36^; *Sema7a*, associated with Multiple Sclerosis^37, 38^ and brain development^39^; and *Zbtb20*, associated with the process of neurodevelopment^40-42^, as well as implicated in major depression disorder^43^. We compared the 12 significant tiles that exhibit age-dependent methylation dynamics to the list of genes dysregulated upon ageing and found only one gene, *Zbtb20*, to be significantly downregulated with age.

### Chromatin accessibility in the cerebral ECs change with age

To study the effect of ageing on chromatin accessibility in the cerebral ECs, we performed the assay for transposase-accessible chromatin sequencing (ATAC-seq) on ECs isolated from male and female mice belonging to 2, 6, 12 and 18 months-old age groups. Across all the samples sequenced, we identified 33254 peaks on an average. In the principal component analysis plot, the samples clustered into two main groups along principal component 3: older mice (12 and 18 months) and younger mice (2 and 6 months). While the principal component 2 (5% of the variance) accounts for the differences between males and females, the principal component 1 cannot be explained by age or any other factors such as sample quality **(Figure 6A)**. Since the second principal component explains 5% variation due to sex, we used a linear model to account for the effects of sex, similar to our analysis of RNA-seq and RRBS data.

**Figure 6.**
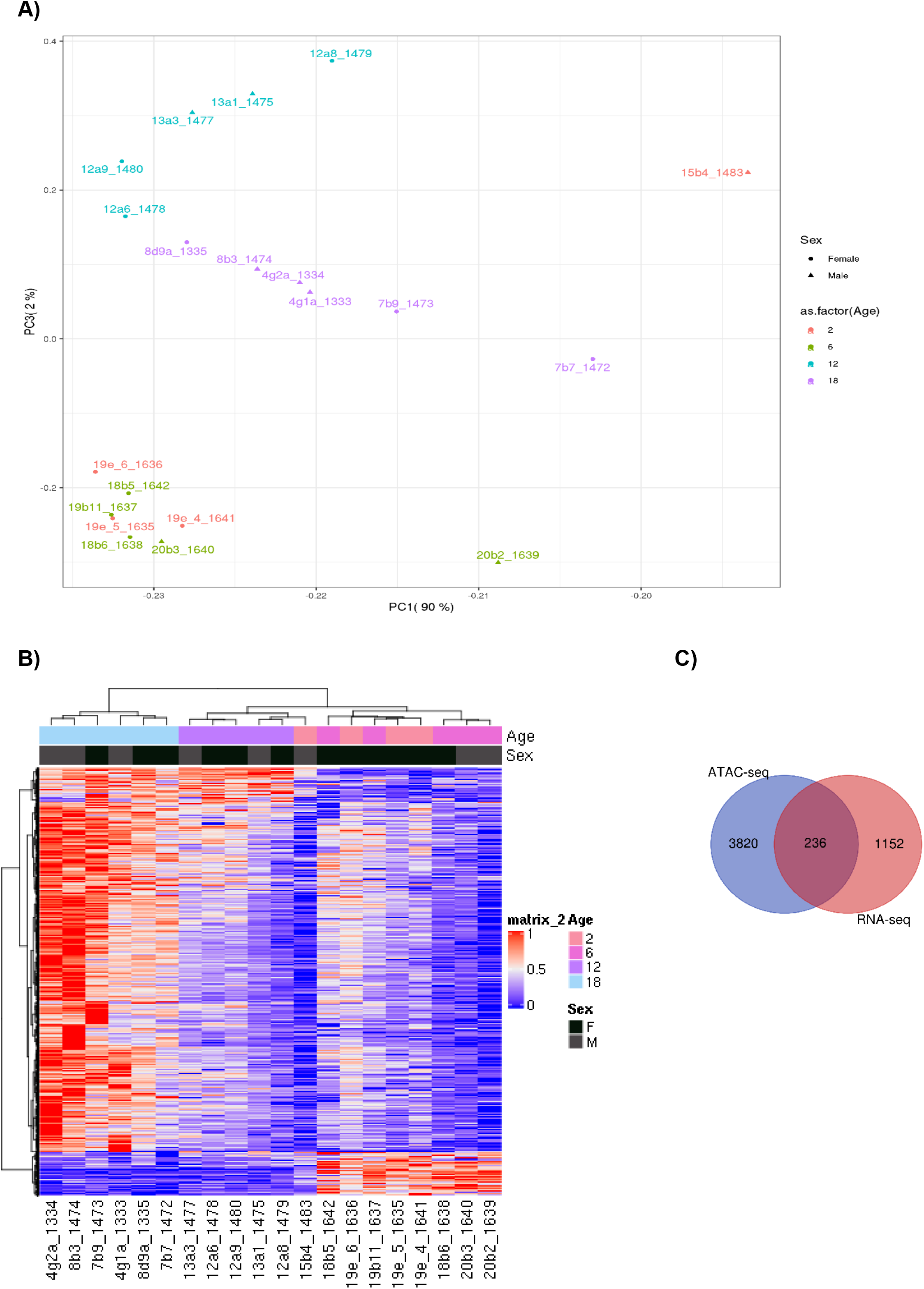
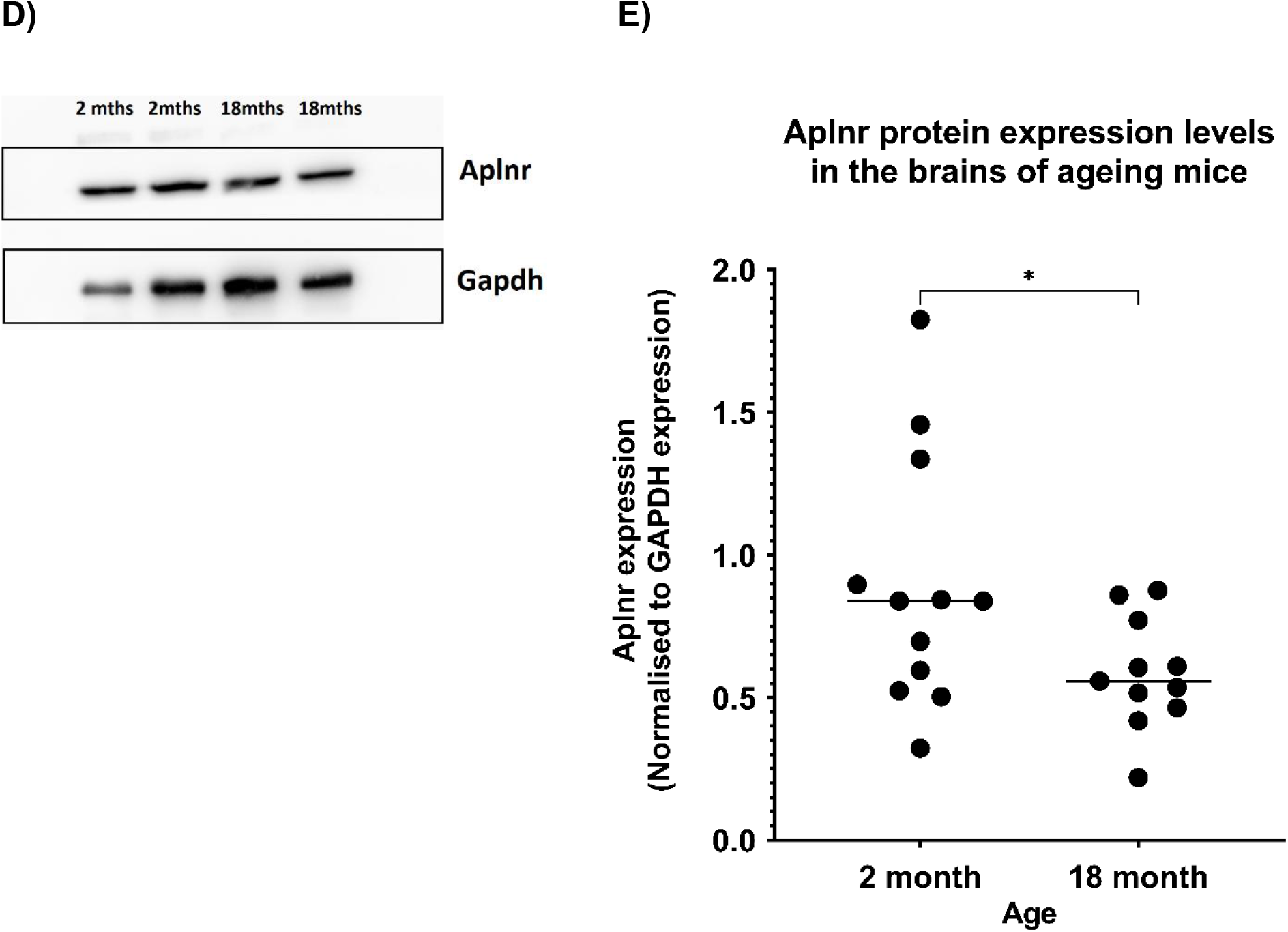
(A) Principal component analysis of samples (ATAC-seq study). The samples cluster into two main groups along principal component 3: older mice (12 and 18 months) and younger mice (2 and 6 months). Principal component 2 accounts for the differences between the sexes, but the principal component 1 cannot be explained by any factor. **(B)** Heatmap showing ATAC signal (normalized per row) in the significant differentially accessible regions (n=11694). **(C)** There are 236 genes that overlap between the list of differentially accessible genes (n=4056) obtained from ATAC-seq analysis and the list of differentially expressed genes (n=1388) based on the RNA-seq analysis. **D)** Immunoblot showing the expression of APLNR and GAPDH proteins in brain lysates prepared from the brains of 2 months-old and 18 months-old mice. **E)** Western blot analysis showed a significant reduction (p value < 0.05) in the expression of apelin receptor in the brains of 18 months-old mice as compared to the 2 months-old mice. The average signal intensity of apelin receptor (normalized to the signal intensity of GAPDH) is 0.89 (sd = 0.44) and 0.58 (sd = 0.19) in the brain lysates prepared from the 2 months- and 18 months-old mice, respectively. Lysates prepared from three mice belonging to both the age-groups were used for western blot studies and the experiment was repeated four times.

Based on the linear model to account for the difference between sex, we extracted 11694 ATAC regions **(Excel S6)** that showed significant change in chromatin accessibility (q-value < 0.05) and plotted a heatmap showing ATAC signal (normalized per row) in these regions (n=11694) **(Figure 6B)** which were annotated afterwards with the closest genes (n=4056). To study the correlation between the age-dependent changes in chromatin accessibility and gene expression, we overlapped the list of differentially accessible genes (n=4056) obtained from ATAC-seq analysis with the list of differentially expressed genes (n=1388) extracted from the RNA-seq analysis. We found 236 genes that overlap between the two gene lists **(Figure 6C)**, including *Aplnr* gene **(Excel S7)**. Interestingly, we found genes of the Apelin signaling pathway (KEGG: mmu04371) such as *Notch3, Gnb4, Mylk* and *Akt3* to be affected by differential upregulation and increased chromatin accessibility. We also compared these 11694 ATAC regions that show significant change in age-related chromatin accessibility to the 12 tiles identified in our methylation study. However, we did not observe any common region or gene. Specifically, while there is a significant age-associated change in chromatin accessibility in the vicinity of the differentially transcribed *Aplnr* gene, we do not observe any change in methylation levels of the region.

### Expression of apelin receptor protein in mouse brain decreases with age

We observed a stark age-associated mRNA downregulation of apelin receptor-encoding gene (*Aplnr*) in cerebral ECs of ageing mice. Interestingly, in the ATAC-seq analysis, we also found an age-dependent reduction in the peak associated with chromatin accessibility of the promoter region of the *Aplnr* gene. We thus investigated the expression levels of apelin receptor protein in ageing mice using western blotting of whole brain lysates prepared from male mice (each in triplicate) of the 2 and 18 months-old age groups. Western blot analysis showed a significant reduction (p-value < 0.05) in the apelin receptor protein levels in 18 months-old mice, as compared to the 2 months-old mice. While the average signal intensity of the apelin receptor (normalized to the signal intensity of the loading control GAPDH) was 0.89 (sd = 0.44) in the brain lysates prepared from the 2 months-old mice, the average signal intensity of apelin receptor (normalized to the signal intensity of GAPDH) was 0.58 (sd = 0.19) in the 18 months-old age group **(Figures 6D,E)**.

## Discussion

In human patients, it has been well-established that age is the severest non-modifiable risk factor for ICH, with incidences of hemorrhages progressively increasing with advancing age^3^. The genetic basis of several human cerebrovascular diseases that affect the integrity of the vasculature in the brain and lead to hemorrhages have been established in recent years. These include the *CCM1,2,3, and KRIT1* genes in cavernous cerebral malformations; *SOX17, CNNM2, KL/STARD13, RBBP8*, and *EDNRA* genes in intracranial aneurysms; *APP, BRI2, CST3, TTR* and *GSN* genes in cerebral amyloid angiopathy and the *COL4A1* and *COL4A2* genes in COL4-related small vessel disease. Additionally, *HTRA1* and *NOTCH3* genes have been implicated in human cerebral autosomal recessive arteriopathy with subcortical infarcts and leukoencephalopathy (CARASIL) and cerebral autosomal dominant arteriopathy with subcortical infarcts and leukoencephalopathy (CADASIL), respectively^44^. These cerebrovascular diseases involve blood vessels of all sizes and lead to the development of microbleeds, intracerebral hemorrhages, or stroke in humans.

We observed that the incidence of cerebral bleedings progressively increases with ageing in mice, reflecting the increased incidence of ICH in elderly human patients. Since endothelial cells form the core component of the BBB in the central nervous system vasculature, ICH is likely caused by molecular defects in EC junction structures leading to the disruption of the blood-brain barrier (BBB) and rupture of blood vessels. Several studies in mice have demonstrated the breakdown of BBB with age, leading to increased BBB permeability^45, 46^. In this work, we profiled the age-dependent changes in transcriptomics, CpG methylation levels, and chromatin accessibility in the cerebral ECs of mice of increasing age. We observed 1388 genes significantly dysregulated with age in the cerebral ECs, with 675 genes downregulated and 713 genes upregulated. Among the major tight junction components, *Cldn5* was downregulated with age in the cerebral ECs of mice. Claudin-5 is an endothelial-specific, highly expressed, vital component of the tight junctions in the cerebral ECs. Interestingly, the *Cldn5* gene is known to be downregulated with age in the cerebral ECs of mice, and aged mice exhibit an increased BBB permeability^46^. Other studies have demonstrated the disruption of BBB and impaired vascular integrity with loss of claudin-5^47, 48^. Further, we observe age-dependent downregulation of *Itm2a*, a highly expressed endothelial-specific transcript, in cerebral ECs. However, the role and function of integral membrane protein 2A in human or murine endothelial cells has not been well characterized. Regarding adherens junction components, Cdh2 was downregulated with age. Transgenic mice with EC-specific deletion of N-Cadherin (*Cdh2*) display impaired vasculature and are embryonic lethal. These mice also show reduced expression of VE-Cadherin in the endothelium, suggesting a critical role of N-Cadherin in vascular morphogenesis^49^.

Cdc42, a small Rho GTPase, is crucial for maintaining the cytoskeletal dynamics of cells by positively regulating actin polymerization and filopodia formation, which further regulates several cellular processes such as cell-cell adhesion, cell division, and cell migration. The role of endothelial Cdc42 in EC-EC adhesion is well studied, with endothelial-specific Cdc42 depletion (Cdc42^*Tie2KO*^ mice) resulting in compromised vasculature, hemorrhages, cerebral vascular malformations^50^ and embryonic lethality^51^. Barry et al. (2015) found that Cdc42 regulates the adhesion between endothelial cells mediated by actin cytoskeletons, physically support the junction complexes^51^. The depletion of endothelial Cdc42 has been reported to disrupt the actin-mediated junctions, thereby compromising the vascular integrity^51^. In our RNA-seq data, we observe a significant age-associated downregulation of *Cdc42*, and several downstream actors involved in the actin reorganization pathway, such as *Cdc42se1, Arf1* and *Arf6*. The downregulation of *Cdc42* in the ageing endothelium might lead to age-associated disruption of the junctional structure at the endothelial cell-cell junction, leading to the age-associated breakdown of the blood-brain barrier and thereby contributing towards increased incidents of cerebral bleedings with age. The actin cytoskeletal dynamics mediated by Cdc42 also regulate Srf, a transcription factor that influences several cellular processes. Interestingly, EC-specific deletion of *Srf* and *Mrtf*, encoding the transcription factors SRF and MRTF-A/-B, at either postnatal or adult ages, induces lethal cerebral hemorrhages in mice^15^. This study was undertaken with the initial hypothesis that an age-dependent increase in bleeding incidents may be due to the age-dependent downregulation of SRF/MRTF function, the dysregulation of SRF/MRTF target genes, and other genes responsible for the maintenance of BBB and vascular integrity in cerebral ECs. However, we did not observe age-dependent significant dysregulation of *Srf* or *Mrtf* **(Figure S10, Table S2)**. However, unaltered mRNA levels do not necessarily correlate with the expression and activity levels of the corresponding proteins. Whether the expression of SRF and MRTF proteins and their activity also remains unchanged in ageing mice’s cerebral ECs remains an open question. However, the observed age-associated downregulation of Cdc42 in ECs may alter SRF activity – another possible pathway that might affect the adhesion between ECs and lead to a compromised blood-brain barrier with age. Any direct roles of endothelial Cdc42 and linked changes in actin dynamics and SRF-regulated target genes remains a possibility contributing to age-associated BBB disruption and therefore warrants further investigation.

This study identifies an age-dependent downregulation of the *Htra1* gene, known to regulate TGF-β signaling, which is necessary to maintain vascular integrity. However, the mechanism of *Htra1* regulating the TGF-β signaling pathway is not well established. While some studies have suggested that Htra1 suppresses TGF-β signaling pathway^52, 53^, others have claimed *Htra1* activates the TGF-β signaling pathway^54^. The detailed molecular pathways underlying the role of HTRA1 in development of CARASIL in human patients are unknown, but HTRA1-mediated dysregulation of TGF-β signaling pathway could lead to vascular impairment in the central nervous system. However, the detailed mechanisms that link HTRA1 and TGF-β in cerebral ECs, and the impact of *Htra1* downregulation on the BBB maintenance needs to be investigated.

We further characterized the changes in DNA methylation with ageing in the cerebral ECs. Previous studies have reported a reduction in global methylation levels, while specific regions may show age-dependent hypermethylation or hypomethylation^55^. This study did not observe substantial global methylation changes in the cerebral ECs upon ageing except for a slight increase in 18 months-old mice. Evidence from recent age-associated methylation studies in mice’s central nervous system suggests that global methylation levels remain largely stable with age in the hippocampus^56^. The methylation levels in the human brain have also been reported to be stable with age^57^. In our study, we observed ageing associated methylation changes for twelve tiles in the vicinity of genes such as Gpr56, *Arid5b, Camta1, Degs2, Sema7a, and Zbtb20* **[Excel Sheet ‘MethTopMarkers’]** previously linked to ageing, neurological development, and diseases. Due to the relatively small number of analyzed individuals, further studies are needed to confirm the identified loci. We then decided to assess the age-dependent changes in chromatin accessibility in the cerebral ECs using ATAC-seq. It is known that the epigenetic changes at the chromatin level have been widely associated with the regulation of gene expression^58^. Our data suggest that the age-dependent chromatin accessibility is associated with gene expression levels in the cerebral ECs. In this study, we observed 11694 peaks undergoing age-dependent change in chromatin accessibility, which were annotated to 4056 genes. Upon comparing these 4056 genes with the 1388 genes found to be dysregulated with age in the RNA-seq dataset, we found that there were 236 genes that are common to both the lists.

One crucial finding of our ATAC-seq analysis was the age-dependent change in chromatin accessibility of the *Aplnr* gene. Since altered chromatin accessibility at the promoter region is associated with altered transcription, the observed age-dependent strong downregulation of *Aplnr* at transcript and protein levels, respectively, represents a very intriguing outcome of the study. Apelin signaling regulates angiogenic sprouting, endothelial tip-cell morphology and sprouting behavior in endothelial cells in zebrafish^22^. Apelin signaling has also been established to positively regulate endothelial cell metabolism as observed by reduced glycolysis in Apelin signaling deficient HUVEC cells^22^. Apelin receptor is known to play a crucial role in positive regulation of vasodilation by heterodimerizing with angiotensin II type 1 receptor (AT1R), leading to its inhibition, thereby negatively regulating the renin-angiotensin system and promoting vasodilation^59, 60^. It has also been reported that Apelin receptor negatively regulates blood pressure in mice. The blood pressure in *Aplnr*^− /−^ mice started to rise around nine months and developed into hypertension when mice attained the age of 12 months^61^. The blood pressure increases with age in both male and female mice, reflecting the pattern observed in humans^62^. Hypertension is an established major risk factor of ICH in humans. These studies suggest that the age-dependent downregulation of *Aplnr* and its ligand *Apln* in cerebral ECs may lead to the activation of the renin-angiotensin system and inhibition of vasodilation, resulting in increased blood pressure in the capillaries. Furthermore, the downregulation of apelinergic axis in mice leads to pathological signs of accelerated ageing, and the infusion of apelin ameliorate age-associated organ impairments, and reduced age-associated cardiovascular pathologies in old mice^61^. Our results suggest an important role of the apelin signaling system in age-associated increase in hypertension and increased brain bleeding incidents. Additionally, the Notch signaling pathway inhibits Apelin signaling. Studies in zebrafish and HUVEC cell line established that the downregulation of Notch leads to an increased expression of Apelin. Conversely, the activation of Notch signaling by treating the HUVECs with Notch ligand DLL4 leads to the reduction in the levels of Apelin. In our RNA-seq and ATAC-seq studies, in addition to the observed effects on the *Aplnr* locus, we also observe an age-dependent upregulation of *Notch3*, consistent with the data that Notch signaling negatively regulates the Apelin signaling pathway.

## Acknowledgements

We thank Dr. Christine Weinl for her valuable advice on the project and providing para-formaldehyde-fixed brain samples of mice, Dr. Angele Breithaupt for her advice on histopathology, Dr. Kristin Bieber and the staff of the Core Facility Flow Cytometry Berg (FCF Berg) at the University Hospital Tübingen for help with the FACS sorting of ECs, Dr. Siegfried Alberti for advice on animal experimentation, and Dr. Kishor Kumar Sivaraj for discussion and advice on animal experimentation, as well as protocols for isolation of endothelial cells.

## Author Contributions

Kshitij Mohan (KM), Alfred Nordheim (AN), Jörn Walter (JW) and Ralf Adams (RA) designed the study and evaluated the results. KM performed histopathology, microscopic analyses and data analyses. KM and Michael Orlich (MO) maintained animal colonies, performed animal experiments and isolated ECs. KM and Gilles Gasparoni (GG) prepared libraries for ATAC-seq and RRBS. GG generated libraries and data of the ATAC-seq study and RRBS. Robert Geffers (RG) was responsible for the RNA-seq study. Abdulrahman Salhab (AS), GG, KM, Steve Hoffmann (SH), RG, JW and AN were responsible for interpretation of data. KM and AN wrote the manuscript.

## References

1. Feigin VL, Krishnamurthi RV, Parmar P, Norrving B, Mensah GA, Bennett DA, et al. Update on the global burden of ischemic and hemorrhagic stroke in 1990-2013: The gbd 2013 study. Neuroepidemiology. 2015;45:161–176

2. van Asch CJ, Luitse MJ, Rinkel GJ, van der Tweel I, Algra A, Klijn CJ. Incidence, case fatality, and functional outcome of intracerebral haemorrhage over time, according to age, sex, and ethnic origin: A systematic review and meta-analysis. The Lancet. Neurology. 2010;9:167–176

3. Jolink WM, Klijn CJ, Brouwers PJ, Kappelle LJ, Vaartjes I. Time trends in incidence, case fatality, and mortality of intracerebral hemorrhage. Neurology. 2015;85:1318–1324

4. Woo D, Broderick JP. Spontaneous intracerebral hemorrhage: Epidemiology and clinical presentation. Neurosurgery clinics of North America. 2002;13:265–279, v

5. Fisher CM. Pathological observations in hypertensive cerebral hemorrhage. Journal of neuropathology and experimental neurology. 1971;30:536–550

6. Harvey A, Montezano AC, Touyz RM. Vascular biology of ageing-implications in hypertension. Journal of molecular and cellular cardiology. 2015;83:112–121

7. Obermeier B, Daneman R, Ransohoff RM. Development, maintenance and disruption of the blood-brain barrier. Nature medicine. 2013;19:1584–1596

8. Abbott NJ, Ronnback L, Hansson E. Astrocyte-endothelial interactions at the blood-brain barrier. Nat Rev Neurosci. 2006;7:41–53

9. Montagne A, Zhao Z, Zlokovic BV. Alzheimer’s disease: A matter of blood-brain barrier dysfunction? The Journal of experimental medicine. 2017;214:3151–3169

10. Stamatovic SM, Johnson AM, Keep RF, Andjelkovic AV. Junctional proteins of the blood-brain barrier: New insights into function and dysfunction. Tissue barriers. 2016;4:e1154641

11. Olson EN, Nordheim A. Linking actin dynamics and gene transcription to drive cellular motile functions. Nat Rev Mol Cell Biol. 2010;11:353–365

12. Weinl C, Riehle H, Park D, Stritt C, Beck S, Huber G, et al. Endothelial srf/mrtf ablation causes vascular disease phenotypes in murine retinae. The Journal of clinical investigation. 2013;123:2193–2206

13. Orlich MM, Diéguez-Hurtado R, Muehlfriedel R, Sothilingam V, Wolburg H, Oender CE, et al. Mural cell srf controls pericyte migration, vessel patterning and blood flow. Circulation research. 2022;131:308–327

14. Deshpande A, Shetty PMV, Frey N, Rangrez AY. Srf: A seriously responsible factor in cardiac development and disease. Journal of Biomedical Science. 2022;29:38

15. Weinl C, Castaneda Vega S, Riehle H, Stritt C, Calaminus C, Wolburg H, et al. Endothelial depletion of murine srf/mrtf provokes intracerebral hemorrhagic stroke. Proc Natl Acad Sci U S A. 2015;112:9914–9919

16. Sindler AL, Delp MD, Reyes R, Wu G, Muller-Delp JM. Effects of ageing and exercise training on enos uncoupling in skeletal muscle resistance arterioles. The Journal of physiology. 2009;587:3885–3897

17. Bird A. DNA methylation patterns and epigenetic memory. Genes Dev. 2002;16:6–21

18. Shih AY, Hyacinth HI, Hartmann DA, Veluw SJv. Rodent models of cerebral microinfarct and microhemorrhage. Stroke. 2018;49:803–810

19. Reuter B, Venus A, Heiler P, Schad L, Ebert A, Hennerici MG, et al. Development of cerebral microbleeds in the app23-transgenic mouse model of cerebral amyloid angiopathy-a 9.4 tesla mri study. Front Aging Neurosci. 2016;8:170

20. Liu S, Grigoryan MM, Vasilevko V, Sumbria RK, Paganini-Hill A, Cribbs DH, et al. Comparative analysis of h&e and prussian blue staining in a mouse model of cerebral microbleeds. The journal of histochemistry and cytochemistry : official journal of the Histochemistry Society. 2014;62:767–773

21. Vanlandewijck M, He L, Mäe MA, Andrae J, Ando K, Del Gaudio F, et al. A molecular atlas of cell types and zonation in the brain vasculature. Nature. 2018;554:475–480

22. Helker CS, Eberlein J, Wilhelm K, Sugino T, Malchow J, Schuermann A, et al. Apelin signaling drives vascular endothelial cells toward a pro-angiogenic state. Elife. 2020;9

23. Goetze JP, Rehfeld JF, Carlsen J, Videbaek R, Andersen CB, Boesgaard S, et al. Apelin: A new plasma marker of cardiopulmonary disease. Regul Pept. 2006;133:134–138

24. Chandra SM, Razavi H, Kim J, Agrawal R, Kundu RK, de Jesus Perez V, et al. Disruption of the apelin-apj system worsens hypoxia-induced pulmonary hypertension. Arteriosclerosis, thrombosis, and vascular biology. 2011;31:814–820

25. Alastalo TP, Li M, Perez Vde J, Pham D, Sawada H, Wang JK, et al. Disruption of pparγ/β-catenin-mediated regulation of apelin impairs bmp-induced mouse and human pulmonary arterial ec survival. The Journal of clinical investigation. 2011;121:3735–3746

26. Strohbach A, Pennewitz M, Glaubitz M, Palankar R, Gross S, Lorenz F, et al. The apelin receptor influences biomechanical and morphological properties of endothelial cells. Journal of cellular physiology. 2018;233:6250–6261

27. Ben-Zvi A, Lacoste B, Kur E, Andreone BJ, Mayshar Y, Yan H, et al. Mfsd2a is critical for the formation and function of the blood-brain barrier. Nature. 2014;509:507–511

28. Ganesh RA, Venkataraman K, Sirdeshmukh R. Gpr56: An adhesion gpcr involved in brain development, neurological disorders and cancer. Brain research. 2020;1747:147055

29. van der Poel M, Ulas T, Mizee MR, Hsiao CC, Miedema SSM, Adelia, et al. Transcriptional profiling of human microglia reveals grey-white matter heterogeneity and multiple sclerosis-associated changes. Nature communications. 2019;10:1139

30. Saare M, Tserel L, Haljasmägi L, Taalberg E, Peet N, Eimre M, et al. Monocytes present age-related changes in phospholipid concentration and decreased energy metabolism. Aging Cell. 2020;19:e13127

31. Mohandas N, Loke YJ, Hopkins S, Mackenzie L, Bennett C, Berkovic SF, et al. Evidence for type-specific DNA methylation patterns in epilepsy: A discordant monozygotic twin approach. Epigenomics. 2019;11:951–968

32. Smith AR, Smith RG, Pishva E, Hannon E, Roubroeks JAY, Burrage J, et al. Parallel profiling of DNA methylation and hydroxymethylation highlights neuropathology-associated epigenetic variation in alzheimer’s disease. Clin Epigenetics. 2019;11:52

33. Vardarajan BN, Barral S, Jaworski J, Beecham GW, Blue E, Tosto G, et al. Whole genome sequencing of caribbean hispanic families with late-onset alzheimer’s disease. Ann Clin Transl Neurol. 2018;5:406–417

34. Sardell RJ, Bailey JN, Courtenay MD, Whitehead P, Laux RA, Adams LD, et al. Whole exome sequencing of extreme age-related macular degeneration phenotypes. Mol Vis. 2016;22:1062–1076

35. Long C, Grueter CE, Song K, Qin S, Qi X, Kong YM, et al. Ataxia and purkinje cell degeneration in mice lacking the camta1 transcription factor. Proc Natl Acad Sci U S A. 2014;111:11521–11526

36. Ohi K, Ursini G, Li M, Shin JH, Ye T, Chen Q, et al. Degs2 polymorphism associated with cognition in schizophrenia is associated with gene expression in brain. Transl Psychiatry. 2015;5:e550

37. Gutiérrez-Franco A, Costa C, Eixarch H, Castillo M, Medina-Rodríguez EM, Bribián A, et al. Differential expression of sema3a and sema7a in a murine model of multiple sclerosis: Implications for a therapeutic design. Clin Immunol. 2016;163:22–33

38. Gutiérrez-Franco A, Eixarch H, Costa C, Gil V, Castillo M, Calvo-Barreiro L, et al. Semaphorin 7a as a potential therapeutic target for multiple sclerosis. Mol Neurobiol. 2017;54:4820–4831

39. Jongbloets BC, Lemstra S, Schellino R, Broekhoven MH, Parkash J, Hellemons AJ, et al. Stage-specific functions of semaphorin7a during adult hippocampal neurogenesis rely on distinct receptors. Nature communications. 2017;8:14666

40. Jones KA, Luo Y, Dukes-Rimsky L, Srivastava DP, Koul-Tewari R, Russell TA, et al. Neurodevelopmental disorder-associated zbtb20 gene variants affect dendritic and synaptic structure. PloS one. 2018;13:e0203760

41. Doeppner TR, Herz J, Bähr M, Tonchev AB, Stoykova A. Zbtb20 regulates developmental neurogenesis in the olfactory bulb and gliogenesis after adult brain injury. Mol Neurobiol. 2019;56:567–582

42. Nagao M, Ogata T, Sawada Y, Gotoh Y. Zbtb20 promotes astrocytogenesis during neocortical development. Nature communications. 2016;7:11102

43. Davies MN, Krause L, Bell JT, Gao F, Ward KJ, Wu H, et al. Hypermethylation in the zbtb20 gene is associated with major depressive disorder. Genome biology. 2014;15:R56

44. Karschnia P, Nishimura S, Louvi A. Cerebrovascular disorders associated with genetic lesions. Cellular and molecular life sciences : CMLS. 2019;76:283–300

45. Stamatovic SM, Martinez-Revollar G, Hu A, Choi J, Keep RF, Andjelkovic AV. Decline in sirtuin-1 expression and activity plays a critical role in blood-brain barrier permeability in aging. Neurobiology of disease. 2019;126:105–116

46. Elahy M, Jackaman C, Mamo JC, Lam V, Dhaliwal SS, Giles C, et al. Blood-brain barrier dysfunction developed during normal aging is associated with inflammation and loss of tight junctions but not with leukocyte recruitment. Immunity & ageing : I & A. 2015;12:2

47. Nitta T, Hata M, Gotoh S, Seo Y, Sasaki H, Hashimoto N, et al. Size-selective loosening of the blood-brain barrier in claudin-5-deficient mice. The Journal of cell biology. 2003;161:653–660

48. Greene C, Hanley N, Campbell M. Claudin-5: Gatekeeper of neurological function. Fluids and Barriers of the CNS. 2019;16:3

49. Luo Y, Radice GL. N-cadherin acts upstream of ve-cadherin in controlling vascular morphogenesis. The Journal of cell biology. 2005;169:29–34

50. Castro M, Laviña B, Ando K, Álvarez-Aznar A, Taha AA, Brakebusch C, et al. Cdc42 deletion elicits cerebral vascular malformations via increased mekk3-dependent klf4 expression. Circulation research. 2019;124:1240–1252

51. Barry DM, Xu K, Meadows SM, Zheng Y, Norden PR, Davis GE, et al. Cdc42 is required for cytoskeletal support of endothelial cell adhesion during blood vessel formation in mice. Development (Cambridge, England). 2015;142:3058–3070

52. Graham JR, Chamberland A, Lin Q, Li XJ, Dai D, Zeng W, et al. Serine protease htra1 antagonizes transforming growth factor-β signaling by cleaving its receptors and loss of htra1 in vivo enhances bone formation. PloS one. 2013;8:e74094–e74094

53. Shiga A, Nozaki H, Yokoseki A, Nihonmatsu M, Kawata H, Kato T, et al. Cerebral small-vessel disease protein htra1 controls the amount of tgf-beta1 via cleavage of protgf-beta1. Human molecular genetics. 2011;20:1800–1810

54. Beaufort N, Scharrer E, Kremmer E, Lux V, Ehrmann M, Huber R, et al. Cerebral small vessel disease-related protease htra1 processes latent tgf-beta binding protein 1 and facilitates tgf-beta signaling. Proc Natl Acad Sci U S A. 2014;111:16496–16501

55. Xiao FH, Kong QP, Perry B, He YH. Progress on the role of DNA methylation in aging and longevity. Briefings in functional genomics. 2016;15:454–459

56. Hadad N, Masser DR, Blanco-Berdugo L, Stanford DR, Freeman WM. Early-life DNA methylation profiles are indicative of age-related transcriptome changes. Epigenetics & Chromatin. 2019;12:58

57. Wagner M, Steinbacher J, Kraus TF, Michalakis S, Hackner B, Pfaffeneder T, et al. Age-dependent levels of 5-methyl-, 5-hydroxymethyl-, and 5-formylcytosine in human and mouse brain tissues. Angewandte Chemie (International ed. in English). 2015;54:12511–12514

58. Benayoun BA, Pollina EA, Brunet A. Epigenetic regulation of ageing: Linking environmental inputs to genomic stability. Nat Rev Mol Cell Biol. 2015;16:593–610

59. Siddiquee K, Hampton J, McAnally D, May L, Smith L. The apelin receptor inhibits the angiotensin ii type 1 receptor via allosteric trans-inhibition. British journal of pharmacology. 2013;168:1104–1117

60. Wu D, He L, Chen L. Apelin/apj system: A promising therapy target for hypertension. Molecular biology reports. 2014;41:6691–6703

61. Rai R, Ghosh AK, Eren M, Mackie AR, Levine DC, Kim SY, et al. Downregulation of the apelinergic axis accelerates aging, whereas its systemic restoration improves the mammalian healthspan. Cell reports. 2017;21:1471–1480

62. Barsha G, Denton KM, Mirabito Colafella KM. Sex- and age-related differences in arterial pressure and albuminuria in mice. Biol Sex Differ. 2016;7:57–57

