## Supplementary Information for "Age-associated changes in endothelial transcriptome and chromatin landscape correlate with elevated risk of hemorrhage"

### Expanded Methods

#### Animal models.

For histological analysis of bleedings in the brains, both wild-type mice and mice having floxed, yet non-recombined *Srf* and *Mrtf* alleles, thereby having functional *Srf* and *Mrtf* alleles (referred to as wild-type phenotype) were used[63]. The mice were generated and housed at the Interfaculty Institute of Cell Biology, Tübingen (Germany). To isolate cerebral endothelial cells, the transgenic Cdh5-mT/H2B-GFP mice were used[64]. Two male transgenic Cdh5-mT/H2B-GFP (heterozygous) mice having a C57BL/6J background were provided by Prof. Dr. Ralf Adams at the Max Planck Institute for Molecular Biomedicine, Münster (Germany). The colony was further expanded at the animal facility in the department of molecular biology at the Interfaculty Institute of Cell Biology, Tübingen (Germany). The mice were kept under 12 hours day/night cycle (daytime from 6 a.m. to 6 p.m) with *ad libitum* access to food and water. The mice were randomly checked for the presence of pathogens and infections every six months. The mice were maintained according to the regulations pertaining to legal animal protection laws, and the experiments performed as part of this project were approved by the Regierungspräsidium Tübingen (Project Nr. ‚Mitteilung nach § 4 Abs. 3 TierSchG, date 18.10.2017). Genotyping of mice was by PCR of ear biopsies. For harvesting brains, the mice were sacrificed using rising concentration of carbon dioxide gas in a gas chamber.

#### Histopathology.

Brains were fixed in 4% PFA solution at 4°C for 72 hours, followed by washing under running cold tap water for 3 hours in a beaker. The fixed brain tissues were further treated with increasing concentrations of isopropanol (50%, 75%, 90%, and 100% v/v) followed by Roti®-Histol. The processed tissues were then embedded in paraffin and coronally sectioned (Leica rotary microtome RM 2155). Four consecutive sections, each 6 µm, were mounted on an adhesive microscope slide (Marienfeld HistoBond®). The sections were dewaxed in Roti®-Histol and treated with decreasing concentrations of ethanol (100%, 96%, 80% and 70% v/v) for rehydration, followed by Hematoxylin-eosin (H&E) staining. The Cover slips were mounted with Entellan as the mounting medium.

**Microscopic analysis.**

Hematoxylin-Eosin stained brain sections were observed under Zeiss Axioplan 2 microscope using an AxioCamHRc camera. To quantify the number of bleedings and microbleedings in each brain, every tenth slide was evaluated, yielding 20 slides per brain for each animal. Images of intact and ruptured blood vessels were taken at same magnification. A blood vessel with extravasation of erythrocytes was counted as a bleeding. If the number of erythrocytes surrounding a ruptured blood vessel was less than or equal to 5, it was scored as a microbleed. The number of bleedings and microbleeds across different age-groups were statistically analyzed using one-way analysis of variance (ANOVA), followed by Tukey's HSD test to compare all the possible pairs and test the statistical significance. The analysis was performed using R and graphs were made using Graphpad Prism version 9.

**Purification of ECs.**

Transgenic Cdh5-mT/H2B-GFP mice were sacrificed by exposure to $CO_2$ gas, followed by cervical dislocation and their brains were harvested. After washing with ice-cold PBS, each brain was cut into 8 sagittal slices and dissociated using the Adult Brain Dissociation Kit (Miltenyi Biotec) according to the manufacturer's instructions. The cell pellet obtained was resuspended in FACS buffer (1X PBS, 2%FCS and 2mM EDTA) to prepare a single-cell suspension from brain. Prior to sorting, 0.05 µg/mL DAPI was added to stain the dead cells and the single-cell suspension was filtered through a 70 µm cell strainer. The Fluorescence Activated Cell Sorting (FACS) was performed on a BD FACS AriaII (BD Sciences) at the FACS Core Facility Berg, Universitätsklinikum Tübingen. Since the transgenic Cdh5-mT/H2B-GFP mice express – specifically in the ECs – red fluorescence in the cell membrane and green fluorescence in the nuclei, cells double positive for tdTomato and GFP were sorted to obtain a pure population of ECs. Single-cell suspensions from brains of littermate wild-type mice lacking the Cdh5-mT H2B-GFP transgene were used as negative controls for FACS.

**RNA isolation and RNA-seq.**

Endothelial cells were directly sorted into RLT Buffer (Qiagen), thereby lysing the cells, and an equal volume 70% ethanol was added to the lysate. The mRNA was isolated and DNase digestion was performed using the RNeasy Micro Kit (Qiagen) according to the manufacturer's instructions. Six animals (3 males and 3 females) belonging to the age-groups of 2, 6, 12, 18 and 24 months (total n = 30) were used for the RNA-seq study. The quality and concentration of the isolated mRNA from each of the thirty samples were assessed using Bioanalyzer RNA 6000 Pico assay (Agilent), with every sample having RIN > 8.3. Full-length cDNA libraries were prepared with the SMART-Seqv4 Ultra Low Input RNA kit (TaKaRa) and the libraries for sequencing were prepared using Nextera XT DNA Library Prep (Illumina) according to the manufacturer's instructions. The libraries were sequenced on Illumina NovaSeq6000 sequencing system at Helmholtz Zentrum für Infektionsforschung, Braunschweig (Germany) and generated 50 bp paired end reads (PE50).

**Assay for Transposase-Accessible Chromatin (ATAC) -seq.**

Cerebral endothelial cells were sorted into FACS buffer (1X PBS, 2%FCS and 2mM EDTA) at 4°C and the assay for transposase-accessible chromatin sequencing (ATAC-seq) was performed according to a protocol adapted from Buenrostro et al[65]. 50,000 endothelial cells were spun down (500g, 5 minutes, 4°C), supernatant was removed and the cell pellet was resuspended in cold DNase inhibiting buffer (1 M KCl, 5 M NaCl, 1 M Tris-HCl, 0.5 M EGTA and 0.5 M Spermidine) containing protease inhibitor cocktail (Roche). 0.1% IGEPAL® CA-630 (Sigma) was added to the suspension followed by gently inverting the tube 3-4 times and incubation on ice for 5 minutes. The nuclei were spun down (500g, 5 minutes, 4°C), supernatant was removed and resuspended in the DNase inhibiting buffer. The nuclei were again centrifuged (500g, 9 minutes, 4°C), the supernatant was removed, and the pellet was resuspended in 50 µL reaction mixture, containing 25 µL 2X TD buffer, 22.5 µL nuclease-free water, and 2.5 µL Tn5 transposase enzyme (Nextera DNA Library Preparation Kit, Illumina, FC-121-1030). The tagmentation reaction was performed at 37 °C for 30 min followed by purification of library using MinElute PCR Purification Kit (Qiagen). The library was eluted in 26 µL elution buffer (Qiagen) and stored at –80°C until amplification. For amplification, 20 µL library was added to 30 µL PCR mix containing 25 µL NEB Next High Fidelity 2X Master Mix (New England Biolabs), 1 µL each Nextera i5 and i7 indexed primers as forward and reverse primers, and 3 µL nuclease-free water. The amplification was carried out in following steps – one cycle of 72°C for 5 minutes and 98°C for 30 seconds, 12 cycles of 98°C for 10 seconds, 63°C for 30 seconds and 72°C for 1 minute and one cycle of 72°C for 5 minutes. The amplified library was cleaned using 0.8x volume (40 µL) of Ampure beads XP (Beckman Coulter) according to the manufacturer's instruction and eluted in 20 µL of 0.1X TE buffer. Six animals (3 males and 3 females) belonging to each of the age-groups of 2, 6, 12 and 18 months (total n = 24) were used for the ATAC-seq study. The quality of the ATAC library was analyzed with Bioanalyzer High-Sensitivity DNA Analysis kit (Agilent) and the concentration of the library was determined using the Qubit HS DNA kit (Life Technologies). The ATAC libraries were sequenced on an Illumina HiSeq2500 sequencing system at the University of Saarland, Saarbrücken and generated approx. 50 million 100 bp paired end reads for each sample.

**Reduced Representation Bisulfite Sequencing.**

100,000 cerebral endothelial cells were sorted in FACS buffer (1X PBS, 2%FCS and 2mM EDTA) at 4°C and centrifuged (500g, 5 minutes at, 4°C). The supernatant was discarded, the cells were snap frozen in liquid nitrogen and stored at –80°C till further processing. For lysing the cells, 200 µL solution

A (25mM EDTA, 75mM NaCl), 200 µL solution B (10 mM EDTA, 10 mM Tris-HCl, 1% SDS) and 10 µL Proteinase K (20 µg/µL) were added to the frozen cell pellet, followed by a brief vortex and incubation at 55°C. Phenol-chloroform-isoamyl (25:24:1) and chloroform-isoamyl (24:1) were used for liquid phase separation of the genomic DNA. Glycogen (20 µg/ µL), 0.1x volume (20 µL) of 3M Sodium Acetate and 2.5x volume (500 µL) of ice-cold 100% ethanol were added and incubated overnight at –20°C for precipitation of genomic DNA. The pellet of genomic DNA obtained was washed with 70% ethanol and dissolved in 40 µL pre-warmed 1x Tris-EDTA (TE buffer) at 45°C for 2 hours. DNA concentration was measured using a Qubit double stranded high sensitivity DNA assay kit (Life Technologies) according to the manufacturer's instructions. Restriction was performed on 26 µL DNA template using 1 µL HaeIII restriction enzyme (New England Biolabs) and 3 µL 10x Cutsmart buffer (New England Biolabs) at 37°C for 18 hours. A-Tailing was performed with 1 µl Klenow Fragment (3′ → 5′exo−, 5 U/µl, NEB) and 1 µl dATP (10 mM, NEB) at 37 °C for 30 min followed by enzyme inactivation at 75 °C for 20 min. Unique molecular identifier adapters (TruSeq Single Index Set B, Illumina) were ligated using 1 µl adapters (10 µM), 0.5 µl T4 Ligase (2,000 U/µl, NEB), 2 µl ATP (10 mM, NEB) and 1 µl Cutsmart buffer (10X, NEB) at 16 °C for 18 hours, followed by enzyme inactivation 65°C for 20 minutes. Bisulfite conversion and subsequent cleanup was performed using EZ-DNA Methylation Gold Kit (Zymo Research) according to the manufacturer's instructions and the bisulfite-converted genomic library was eluted in 24 µL nuclease-free water. The library was amplified by polymerase chain reaction using 0.6 µl each of primers (10 µM, Primer i5: AATGATACGGCGACCACCGAGATCTACAC, Primer i7: CAAGCAGAAGACGGCATACGAGAT), 0.6 µl Hot Start Taq (5 U/µl, Qiagen), 3 µl Hotstar PCR Buffer (10X, Qiagen), 1.2 µl $MgCl_2$ (25 mM) and 2 µl dNTPs (10 mM) in a 30 µl reaction with 95°C for 15 min, 20 cycles of 95°C for 40 s, 58°C for 1 minute, 72°C for 1 min, 72°C for 12 min and hold at 4°C. The amplified RRBS library was cleaned using 0.8x volume (40 µL) of Ampure beads XP (Beckman Coulter) according to the manufacturer's instruction and eluted in 20 µL of 0.1X TE buffer. Six animals (3 males and 3 females) belonging to each of the age-groups of 2, 6, 12, and 18 months (total n = 24) were used for RRBS. The RRBS library was analyzed with Bioanalyzer High-Sensitivity DNA Analysis kit (Agilent), and the concentration was determined using the Qubit HS DNA kit (Life Technologies). The sequencing was performed on Illumina HiSeq2500 sequencing system at the University of Saarland, Saarbrücken, and generated approx. 50 Million 100 bp single-end reads per sample.

**RNA-seq data processing.**

The preliminary quality control checks on the raw RNA-seq data were performed using FASTQC (v0.11.4) (https://www.bioinformatics.babraham.ac.uk/projects/fastqc/). The 3' adapter sequence (CTGTCTCTTATACACATCTGACGCTGCCGACGA) were trimmed using Cutadapt[66] (v1.15) and quality control checks were again performed on the trimmed reads with FASTQC. The trimmed reads were then aligned to the mouse genome (GRCm38/mm10) using STAR aligner (v2.5.2b)[67]. The parameter *quantMode* was set to *GeneCounts* for the calculation of counts per gene. A comprehensive quality report for all the samples was generated with MultiQC[68] (v1.7) (https://multiqc.info/). The counts obtained after STAR alignment was used to study differential expression analysis using DESeq2[69] (v1.24.0). Linear regression analysis was performed on TPM (transcripts per million) values, calculated by normalizing the raw read counts to the gene length and sequencing depth in each sample, to study the age-associated changes in gene expression. A gene was considered to be significantly dysregulated with age if it had an adjusted p-value less than 0.05 (p.value.age.fdr < 0.05) in the linear regression analysis.

**ATAC-seq data processing**.

The adapter sequence and 3' ends with base quality (PHRED score) of less than 20 in the FASTQ files obtained from the sequencing were trimmed with Trim Galore (v0.4.2) software. Following adapter trimming and removing low quality nucleotides, the FASTQ files were mapped to the mouse reference genome (GRCm38/mm10) using the GEM[70] mapper. Duplicated reads found after alignment with GEM mapper were annotated with Picard tools (v1.115) (http://broadinstitute.github.io/picard). MACS2[71] (v2.1.0) was used to call Nucleosome Depleted Regions (NDRs) after downsampling the reads into similar number to avoid any bias in the downstream analyses due to sequencing depth. The parameters used in the MACS2 were: –shift -100, –extsize 200, –nomodel and –keep-dup all. Differentially accessible regions were calculated using a linear model accounting for gender. Peaks which overlap with the black list regions defined by ENCODE (https://doi.org/10.1038/s41598-019-45839-z) were excluded from the analysis.

**RRBS data processing.**

Sequencing reads were trimmed using the Trim Galore (v0.4.2) software (http://www.bioinformatics.babraham.ac.uk/projects/trim_galore/) to remove the adapter contamination and the 3' ends with base quality (PHRED score) of less than 20. The trimmed reads were then aligned to the mouse reference genome (GRCm38/mm10) using the BWA[72] (v0.6.2) wrapper methylCtools[73] (v0.9.2). Samtools[74] (v1.3) and Picard tools (v1.115) (http://broadinstitute.github.io/picard) were used for converting, merging, and indexing of the alignment files. SNP aware realignment to identify single nucleotide polymorphisms (SNPs) for accurate identification of methylated cytosines and methylation calls were performed with the bisulfite SNP calling software Bis-SNP[75]. MethylKit[76] (v1.3.1) was used for tiling (1kb tiles, min 3 CpGs per tile, each CpG with minimum coverage of 5) . Association of DNA methylation and ageing was calculated using a linear regression model with gender and coverage as covariates. Obtained p-values were corrected for multiple testing using Benjamini Hochberg correction.

**Data access and codes.**

The raw and processed sequencing RNA-seq, ATAC-seq and RRBS files have been deposited to GEO under the accession number GSE218649. Codes used in the analysis of RNA-seq, ATAC-seq and RRBS can be provided on request.

**Western blotting.**

Brain tissue was lysed in radioimmunoprecipitation assay buffer [50 mM Tris·HCl, pH 8, 150 mM NaCl, 1% IGEPAL CA-630, 0.5% Sodiumdeoxycholate, 0.1% SDS, 1 mM EDTA, protease inhibitor (cOmplete™ Mini, Roche)] using mechanical homogenization (Polytron, 2 x 20 seconds), and protein concentration was determined using Bradford assay. Proteins were separated by SDS/PAGE (90V for 30 minutes, 110V for 60 minutes, 130V for 30 mins) and transferred onto 0.45 µm polyvinylidene difluoride transfer membrane (Merck) using Bio-Rad transfer system (0.1 Ampere, 60 minutes). For immunoblotting, the membranes were blocked in 5% nonfat dried milk in TBST for 60 minutes, washed three times with 1X TBST and incubated in primary antibodies (anti-Aplnr, 1:300, Invitrogen #711101 and anti-GAPDH, 1:50,000, Acris #ACR001P) at 4°C for 18 hours. Incubation in corresponding HRP-conjugated secondary antibodies (HRP-linked anti-rabbit IgG, 1:10,000 and HRP-linked anti-mouse, 1:10,000 , GE Healthcare) was done for 60 minutes at room temperature. Membranes were imaged in the Fusion SL documentation system after adding chemiluminescent HRP substrate (Milipore) and quantified using Fusion FX software (Vilber). Gapdh was used as loading control and the expression level of Aplnr was normalized to the expression levels of Gapdh in each sample. The brain lysates for western blot studies were prepared from (n=3 mice) belonging to each of the age-groups and the western blot experiment was repeated four times, thereby using 12 data points from each age-group.

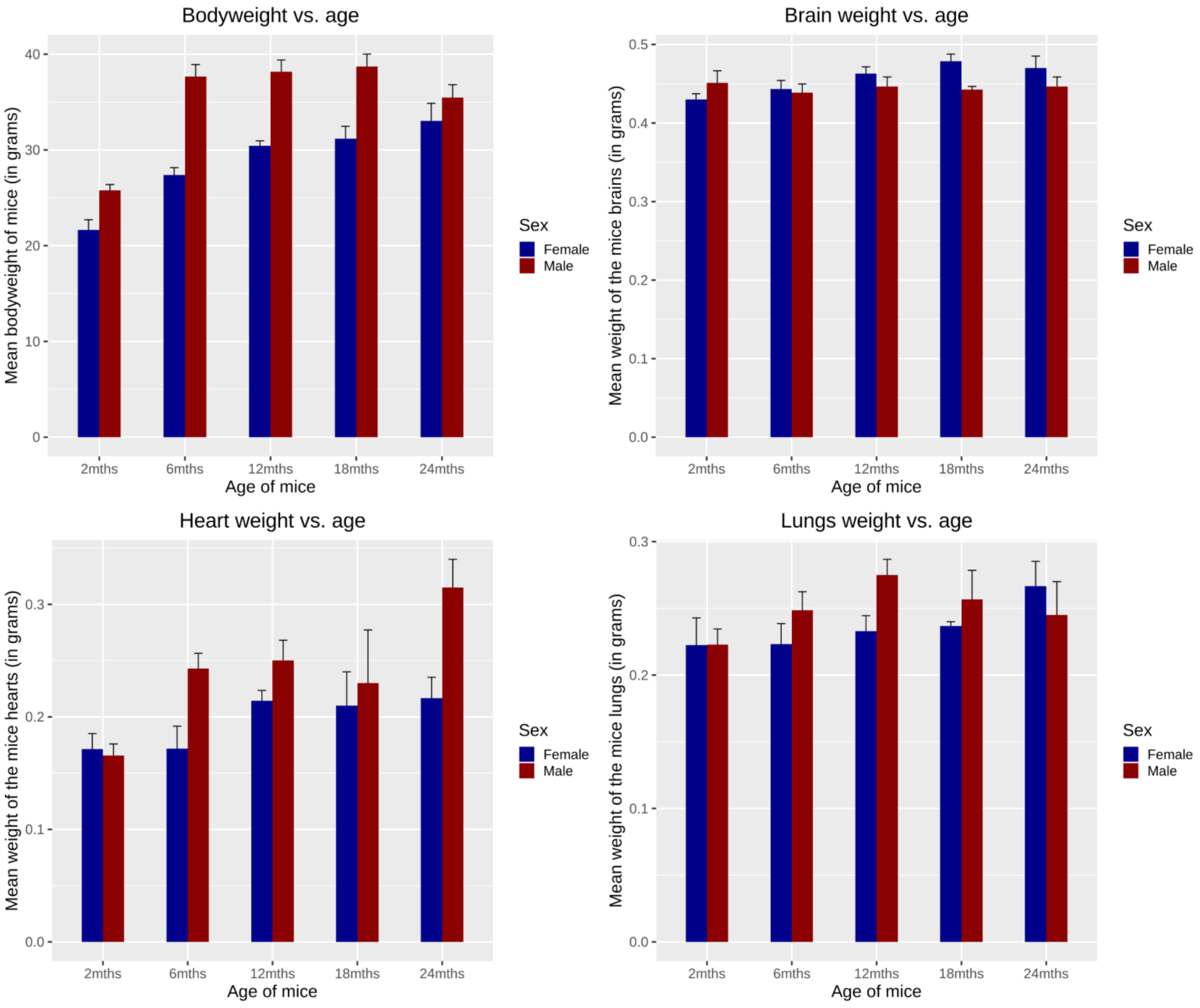


**Figure S1 | Weight of body, brain, heart and lungs of mice as a function of age.** The body weight of mice increases in the 6 months-old group as compared to the 2 months-old mice; however, there is no further significant change in the body weight of mice after the age of 6 months. Similarly, ageing has no effect on the weights of brains, lungs and heart in the mice as the weights of these organs do not change significantly with age. Error bars represent mean mean ± se (standard error of the mean). The mean weights and standard deviation (sd) of each group is summarized in Table 3.1. (n = 6 males and 6 females for the 2 months, 6 months, 12 months and 18 months-old cohorts ; n= 3 males and 3 females for the 24 months-old cohort).

A)

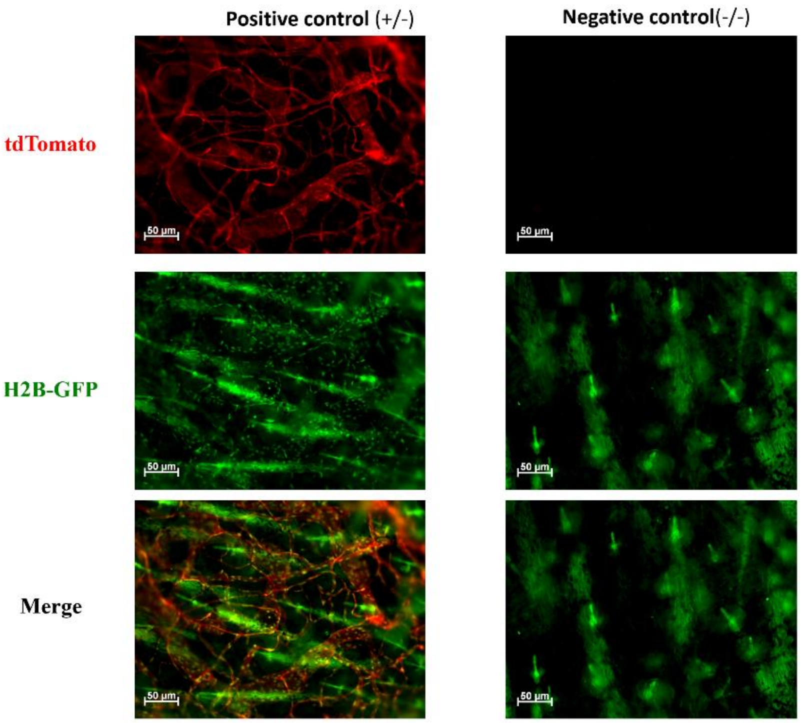


B)

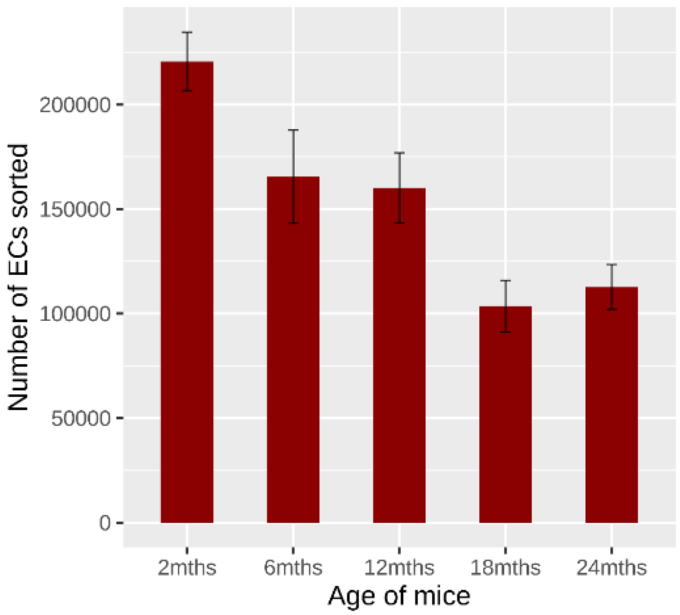


**Figure S2 | ECs isolated from Cdh5-mT/H2B-GFP mice**.

A) A section of ear vasculature from the Cdh5-mT/H2B-GFP mice under the microscope shows blood vessels staining with red fluorescence (membrane targeted Tomato) and the nuclei exhibit green fluorescence (H2B-GFP).

B) The yield of ECs sorted from the brain of older mice is less than the yield from younger mice. The average numbers of ECs sorted from each brain of a 2 months-, 6 months-, 12 months-, 18 months- and 24 months-old mice were 220559 (sd = 52741), 165505 (sd = 77175), 160108 (sd = 70983), 103516 (sd = 49352) and 112745 (sd =26225), respectively. Error bars represent mean ± se (standard error of the mean). An interesting observation of the study has been reduction in the yield of cerebral ECs with ageing, which can be attributed to technical reasons. Much larger amounts of myelin and cellular debris are obtained during the process of preparing single-cell suspension from the brains of 18 and 24 months-old mice (visual observation, not quantified). A plausible explanation for reduced yield of cerebral ECs from older mice is loss of ECs during the step of myelin and cellular debris removal.

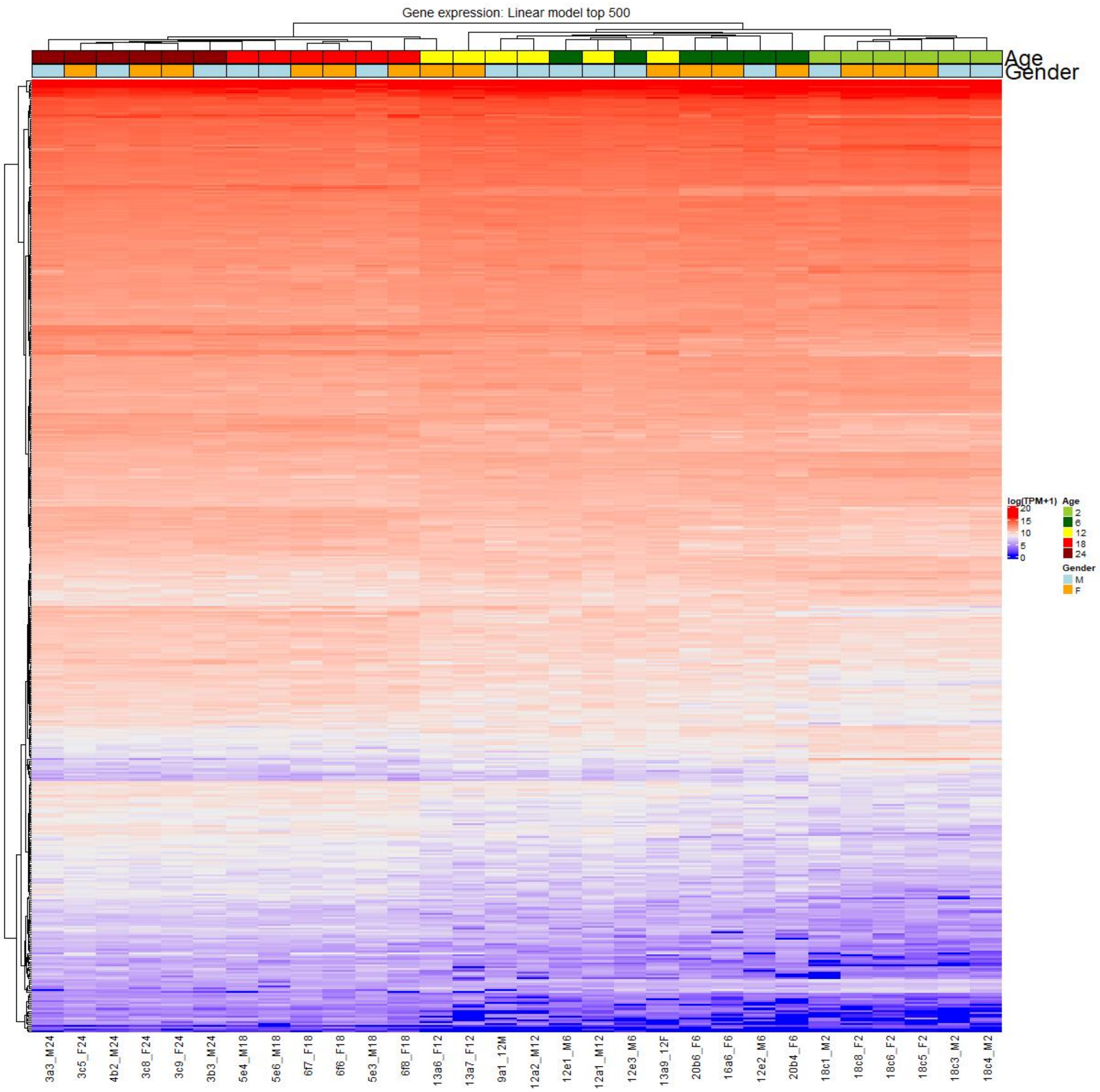


**Figure S3 | Heatmap of top 500 dysregulated genes.** The heatmap of top 500 significantly dysregulated genes in the linear regression model indicate that the expression level of these genes is different across the age-groups. The samples from a particular age-group show similar expression profile and cluster together, suggesting a strong age-associated expression pattern of these genes.

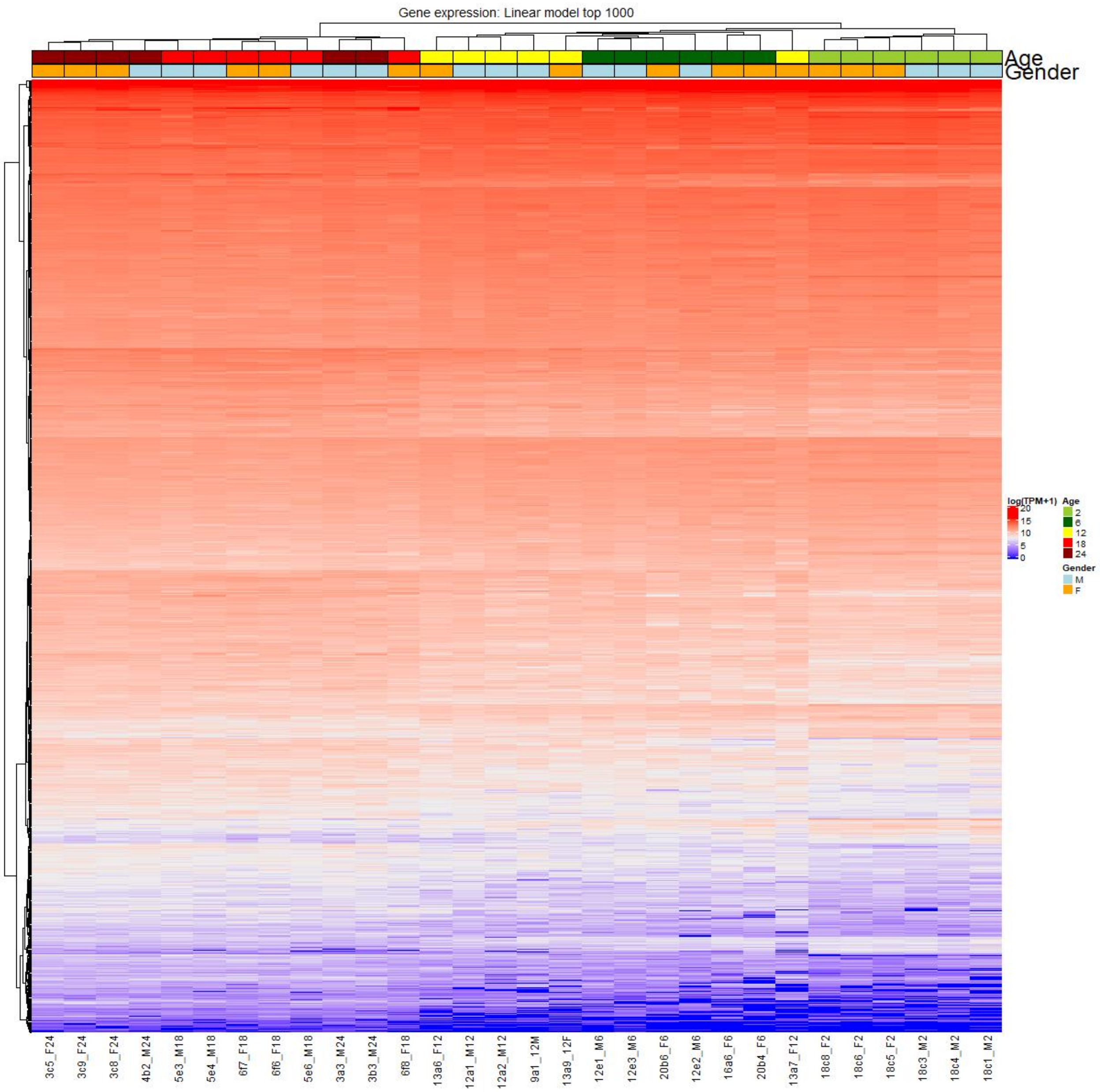


**Figure S4 | Heatmap of top 1000 dysregulated genes.** The heatmap of top 1000 significantly dysregulated genes in the linear regression model also suggests the difference in expression level of these genes across the age-groups. The samples from a particular age-group show similar expression profile and cluster together, suggesting a strong age-associated expression pattern of these genes.

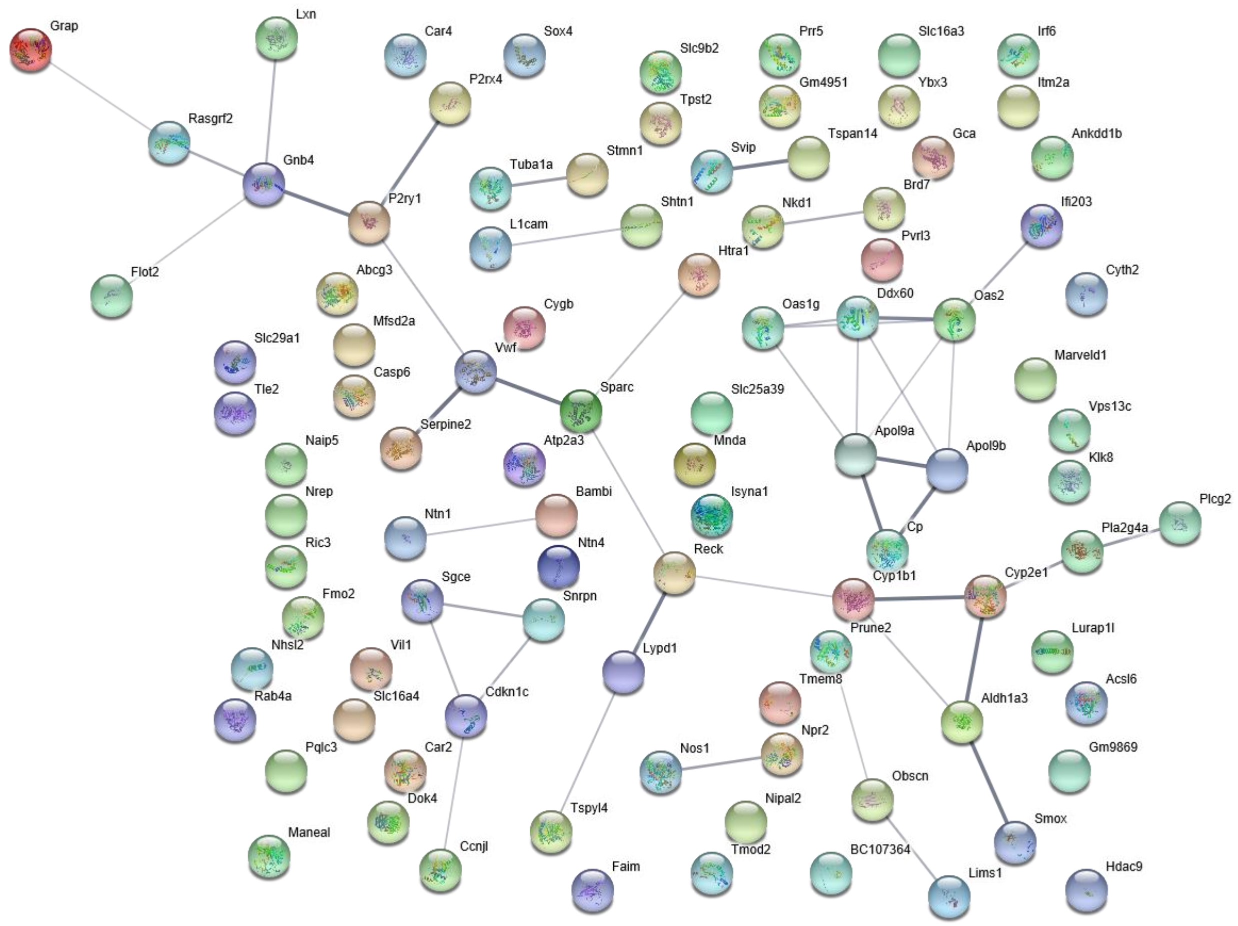


**Figure S5 | Gene enrichment analysis of top 100 age-dependent dysregulated genes based on the RNA-seq linear regression analysis.** The figure represents the enriched processes and the genes associated with those processes. The excel sheet 'Top 100 genes enrichment analysis' depicts all the enriched pathways, their GO numbers and the associated genes.

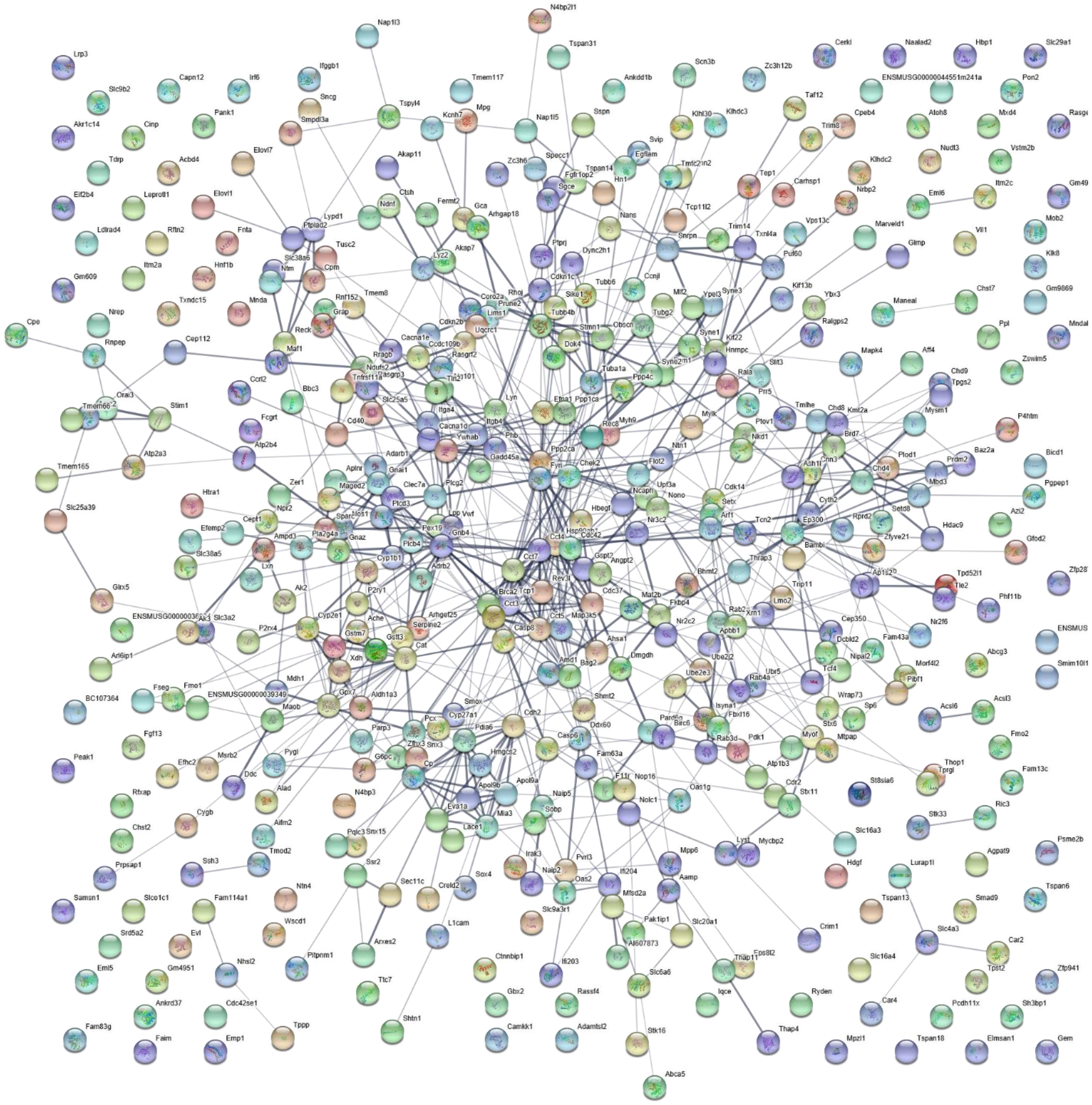

**Figure S6 | Gene enrichment analysis of top 500 age-dependent dysregulated genes based on the RNA-seq linear regression analysis.** The figure represents the enriched processes and the genes associated with those processes. The excel sheet ‘Top 500 genes enrichment analysis’ depicts all the enriched pathways, their GO numbers and the associated genes.

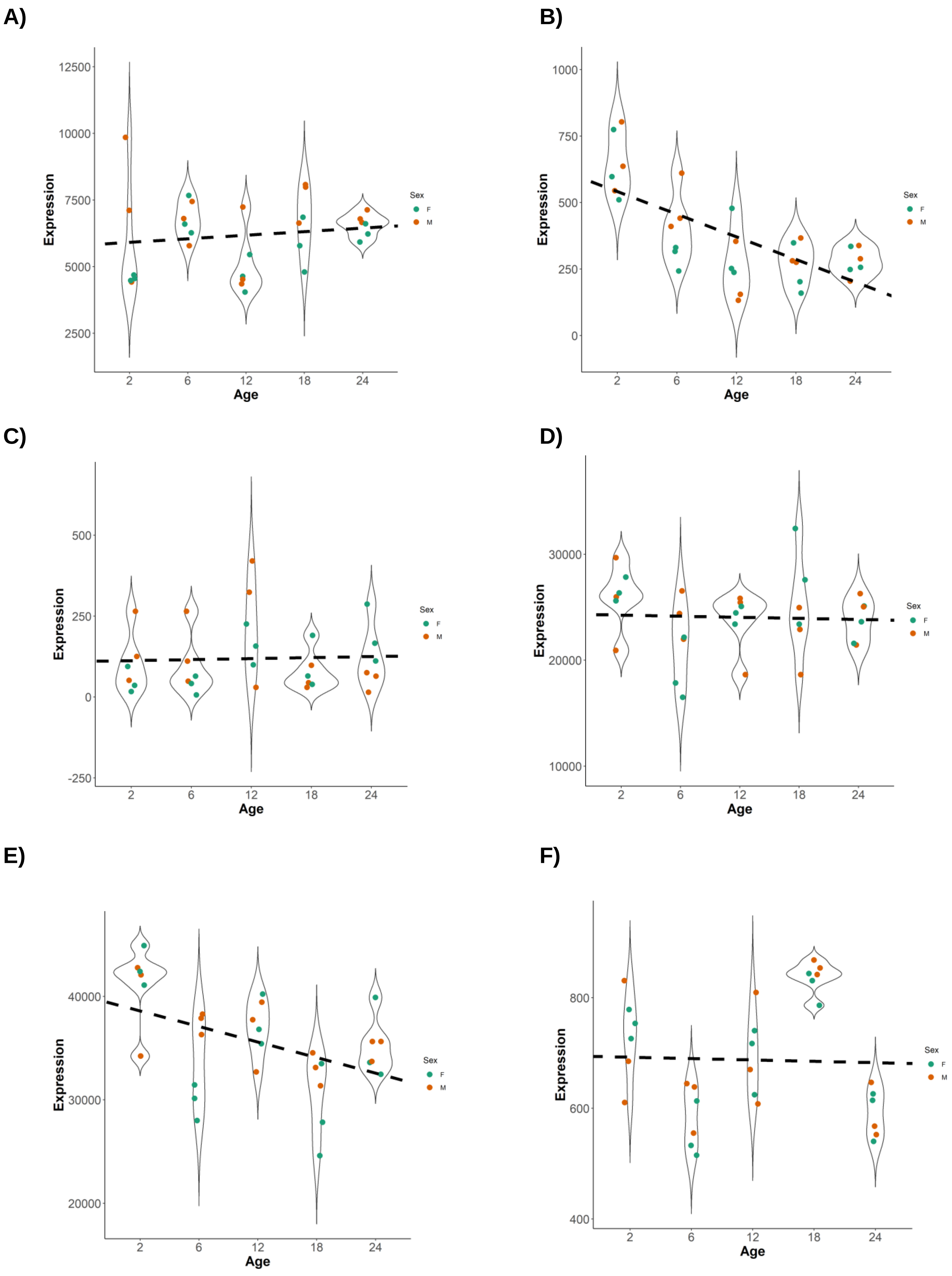
A)
B)
C)
D)
E)
F)
Expression
Age
Sex
F
M
2
6
12
18
24
12500
10000
7500
5000
2500
1000
750
500
250
0
500
250
0
-250
30000
20000
10000
40000
30000
20000
800
600
400

**Figure S7 | Expression of genes encoding key components of adherens junction across age.** The **A)** *Cdh5* gene, which encodes VE-Cadherin, was not dysregulated with age. However, the expression of B) Cdh2 gene encoding for neuronal cadherin showed a significant age-dependent downregulation, with correlation coefficient of 0.45 (adjusted p-value = 0.006). Other major components of the adherens junction such as **C)** Cdh1, **D)** Ctnna1, **E)** Ctnnb1, and **F)** Ctnnd1 were not dysregulated with age (adjusted p-values > 0.1).

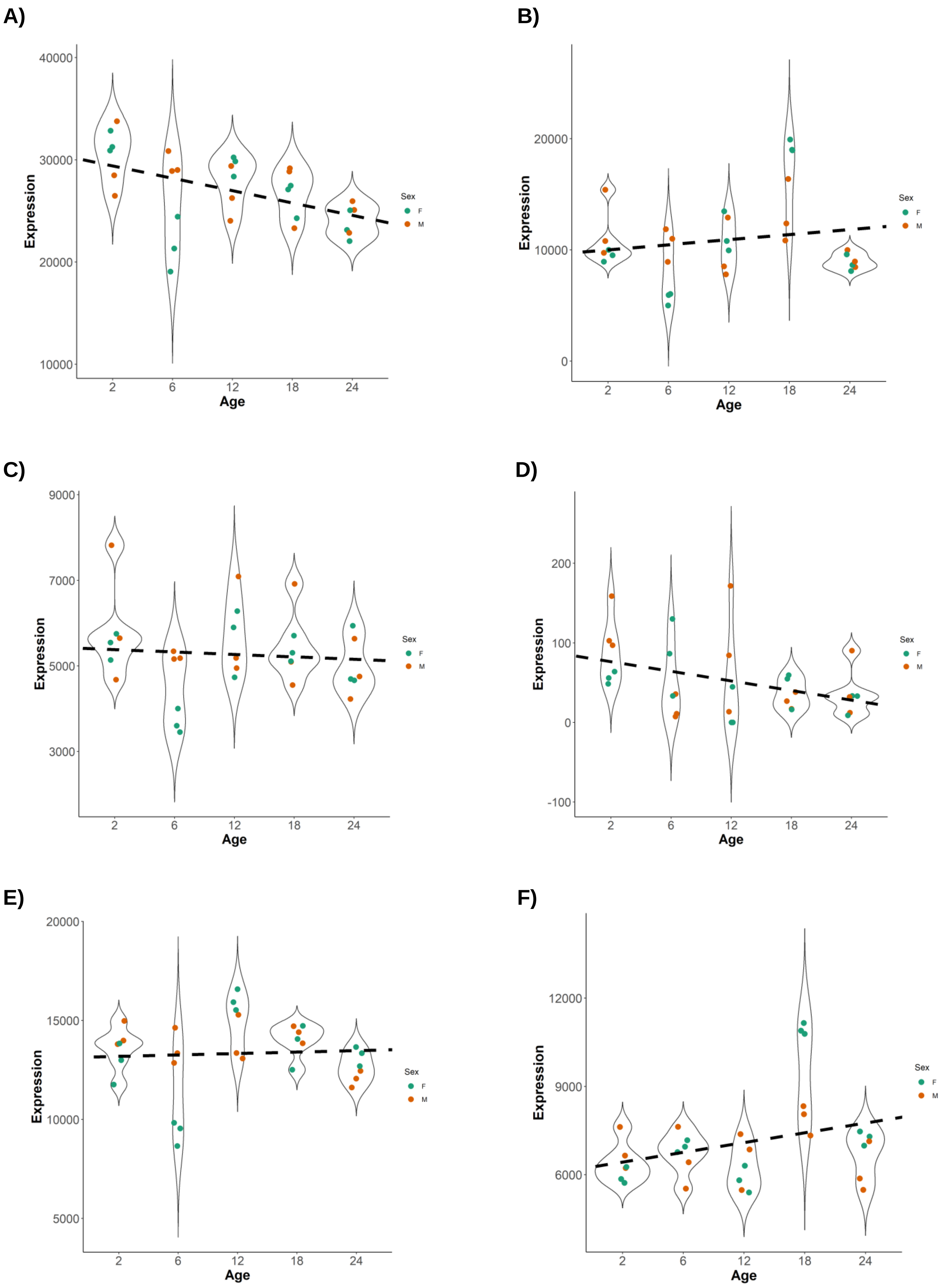

A)
B)
C)
D)
E)
F)
Expression
Age
Sex
F
M
40000
30000
20000
10000
20000
10000
0
9000
7000
5000
3000
200
100
0
-100
20000
15000
10000
5000
12000
9000
6000
2
6
12
18
24

**Figure S8 | Expression of genes encoding key components of tight junction across age.** Several key genes encoding tight junction complex such as **A)** *Jam2*, **B)** *Tjp1*, **C)** *Tjp2*, **D)** *Tjp3* **E)** *Cgn,* and **F)** *Afdn* were not dysregulated with age.

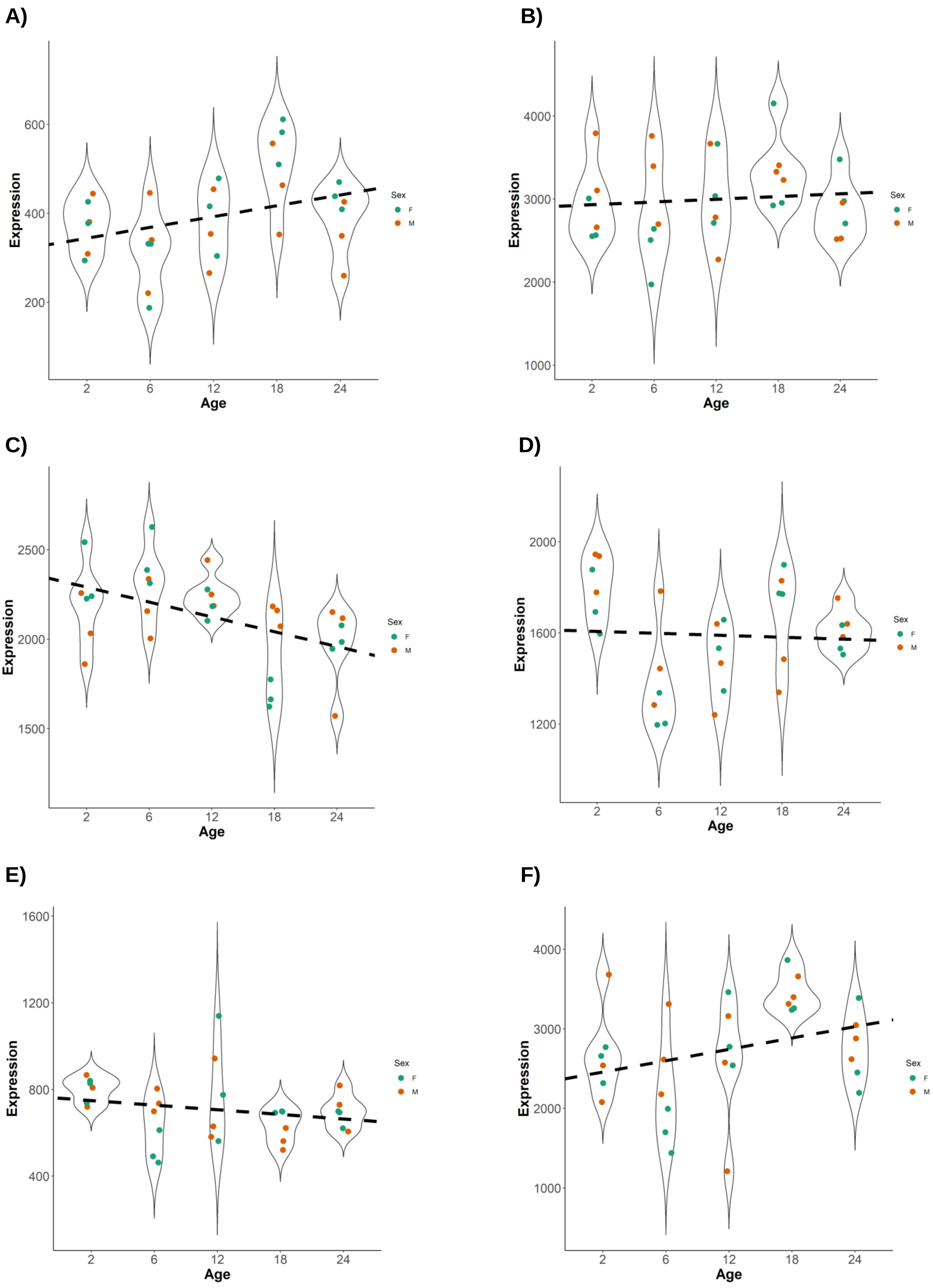
A)
B)
C)
D)
E)
F)
Expression
Age
Sex
F
M
2
6
12
18
24

**Figure S9 | Expression of genes encoding key components of major scaffolding proteins of the blood-brain barrier across age.** We did not observe any significant age-dependent dysregulation of major scaffolding proteins present at the BBB such as **A)** *Magi1*, **B)** *Magi3*, **C)** *Mpp1*, **D)** *Mpp5*, **E)***Mpp7*, and **F)** *Pard3*.

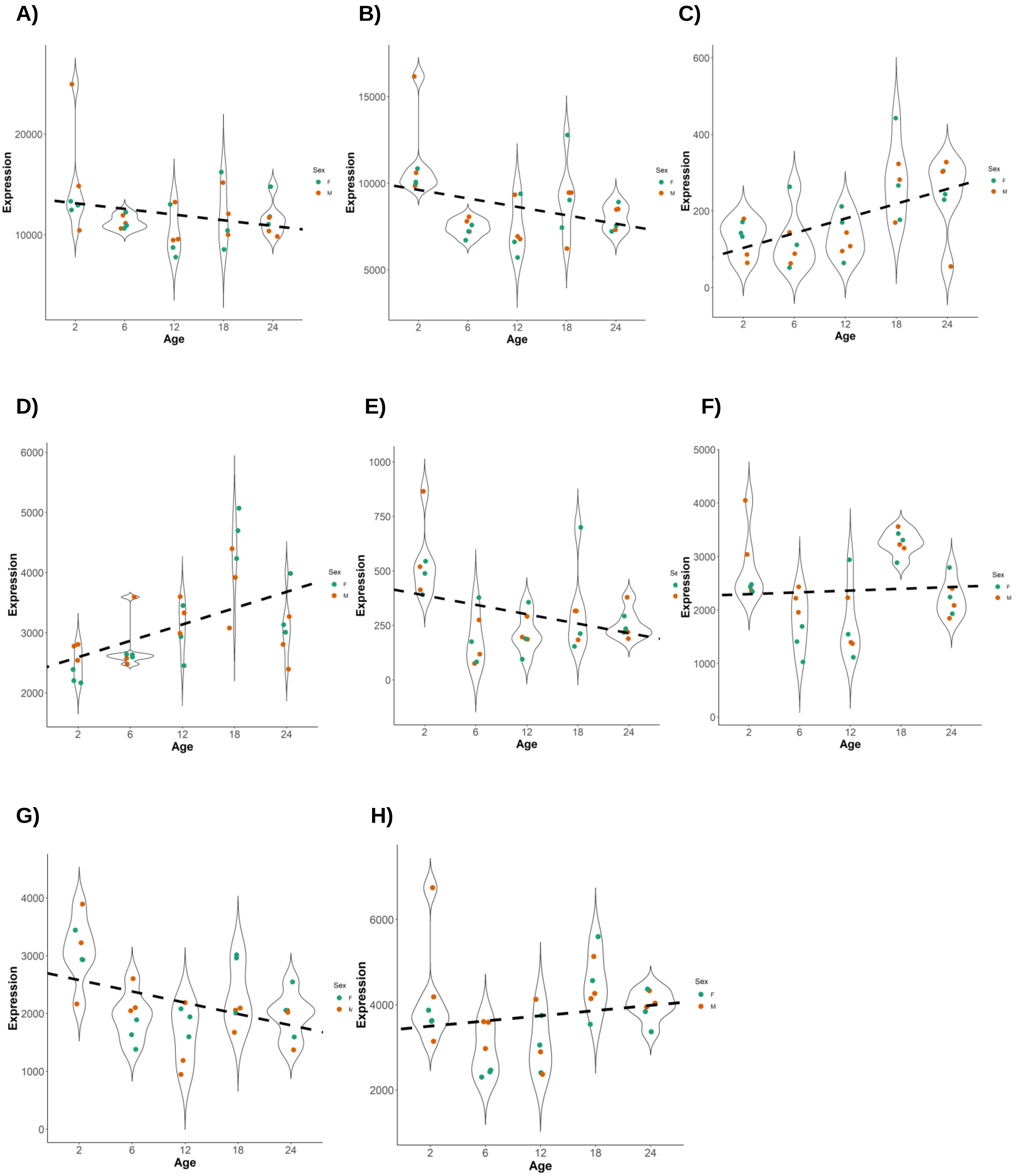

A)
B)
C)
D)
E)
F)
G)
H)
Expression
Age
Sex
F
M
2
6
12
18
24
10000
20000
5000
15000
0
200
400
600
2000
3000
4000
5000
6000
250
500
750
1000

**Figure S10 | Expression of genes encoding key components of basement membrane across age.** The expression levels of transcripts of several major components of basement membrane and extracellular matrix proteins implicated in the maintenance of the BBB were found to be unchanged with age. **A)***Col4a1*, **B)***Col4a2*, **C)***Lama2*, **D)***Lama5*, **E)***Lamb1*, **F)***Lamc1*, **G)***Nid1* and **H)***Nid2* do not exhibit any significant age-dependent dysregulation in the linear regression analysis of the RNA-seq dataset (adjusted p-values > 0.1).

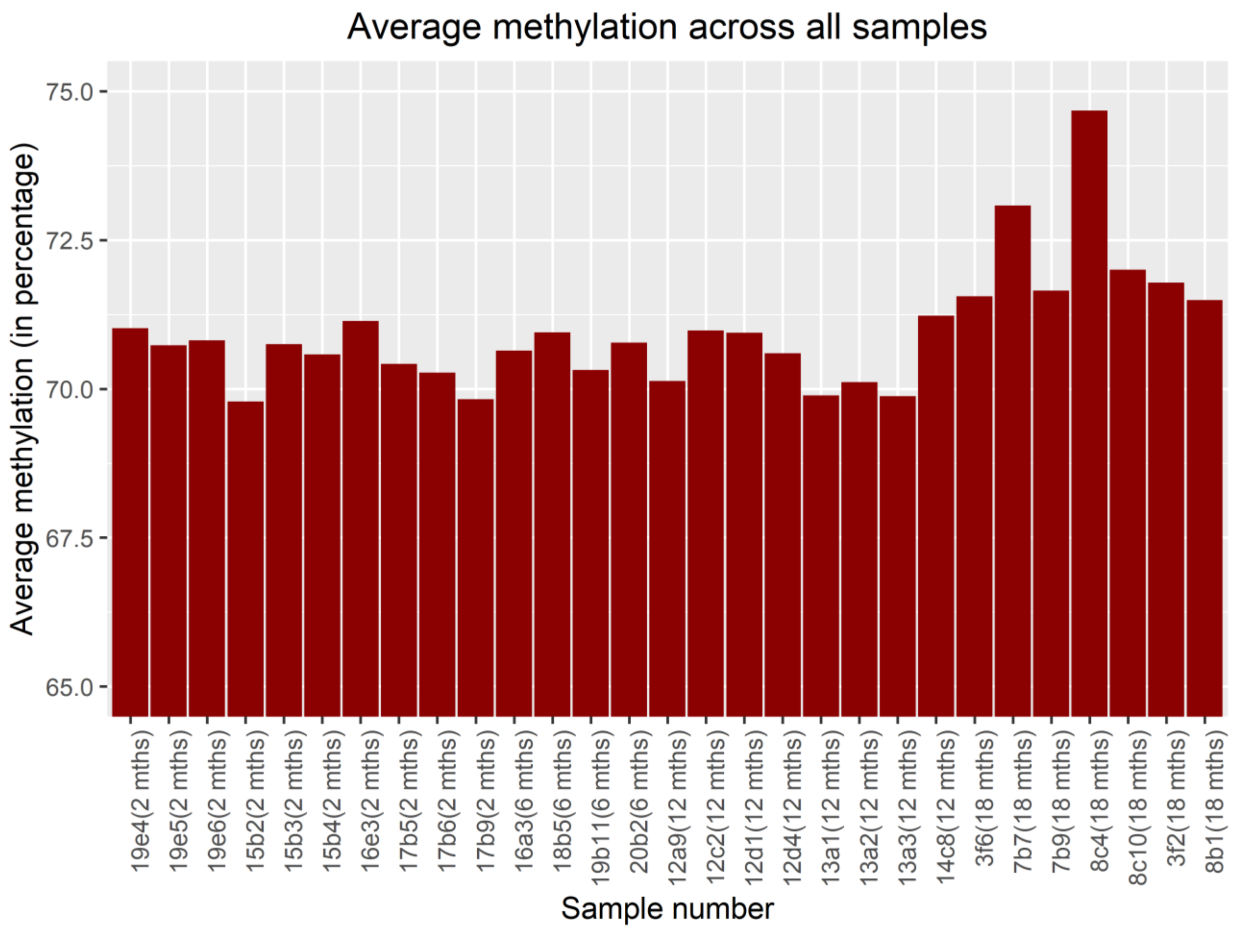


**Figure S9 | Average methylation across all the samples**. The bar plot shows average methylation (in percentage) across all the samples considered for RRBS analysis. The x-axis indicates the name of the mouse and its age in parentheses.

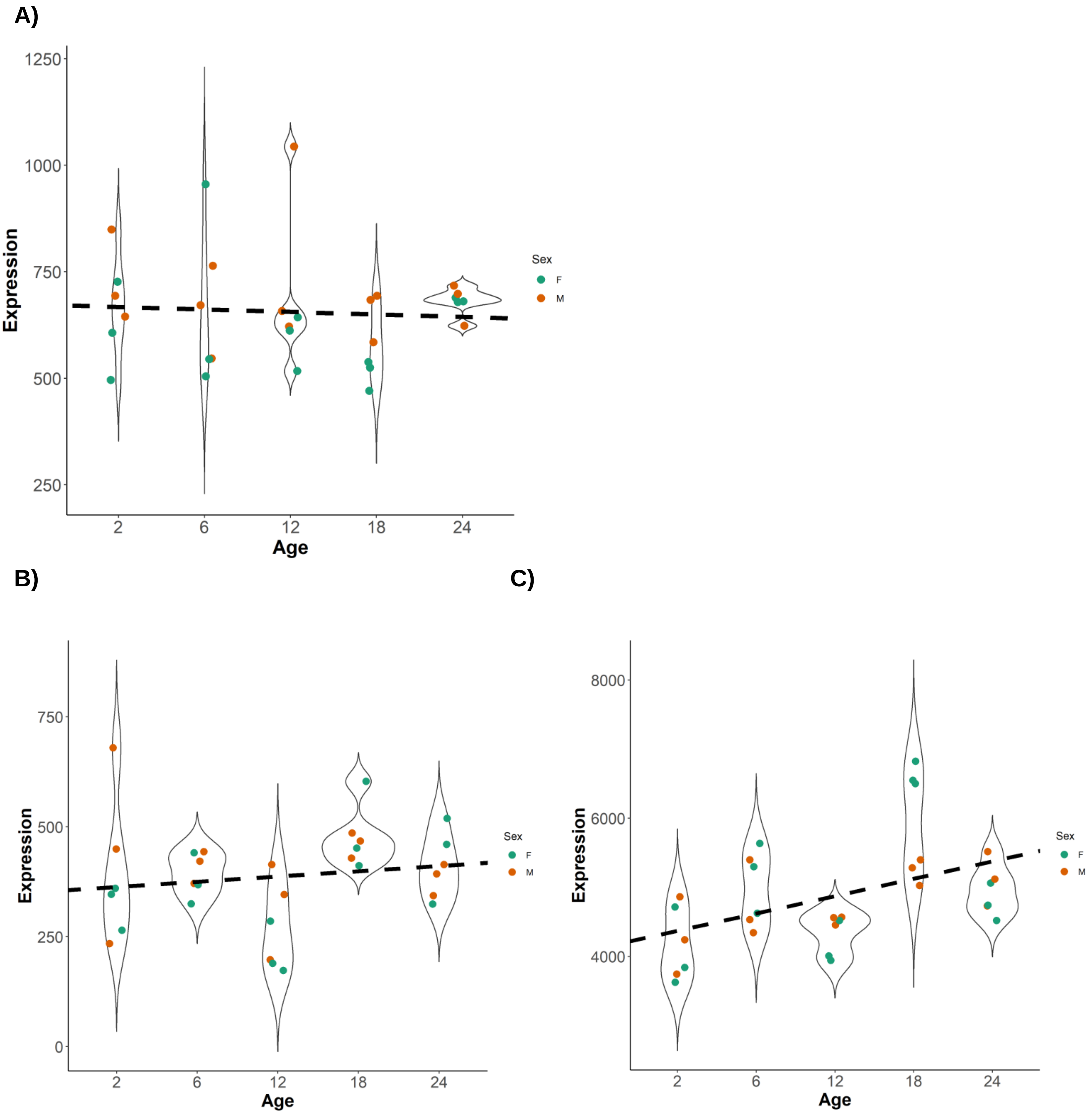

A)
B)
C)
Expression
Age
Sex
F
M
1250
1000
750
500
250
0
8000
6000
4000
2
6
12
18
24

**Figure S10 | The expression level of *Srf*, *Mrtfa* and *Mrtfb* with age**. Linear regression analysis of RNA-seq dataset suggests that the expression of **(A)** *Srf* (adjusted p-value = 0.88), **(B)** *Mrtfa* (adjusted p-value = 0.69) and **(C)** *Mrtfb* (adjusted p-value = 0.15) does not change significantly with age. The linear regression analysis performed on the RNA-seq dataset includes the males and females but corrects for the effects of sex by taking the differences in expression between males and females as covariates. The linear regression analysis was performed on the TPM (transcripts per million) values calculated for all the transcripts in each sample.

| Age | Sex | Body weight (gram) Mean ± sd | Brain weight (gram) Mean ± sd | Heart weight (gram) Mean ± sd | Lungs weight (gram) Mean ± sd |
|---|---|---|---|---|---|
| 2 months | Male | 25.76 ± 1.87 | 0.451 ± 0.046 | 0.165 ± 0.026 | 0.222 ± 0.031 |
| | Female | 21.64 ± 3.20 | 0.43 ± 0.217 | 0.171 ± 0.039 | 0.222 ± 0.057 |
| 6 months | Male | 37.66 ± 3.31 | 0.438 ± 0.029 | 0.242 ± 0.036 | 0.248 ± 0.034 |
| | Female | 27.39 ± 1.86 | 0.443 ± 0.026 | 0.171 ± 0.049 | 0.223 ± 0.037 |
| 12 months | Male | 38.18 ± 2.99 | 0.446 ± 0.029 | 0.25 ± 0.044 | 0.275 ± 0.028 |
| | Female | 30.41 ± 1.41 | 0.462 ± 0.022 | 0.214 ± 0.024 | 0.232 ± 0.030 |
| 18 months | Male | 38.71 ± 3.68 | 0.442 ± 0.011 | 0.23 ± 0.081 | 0.256 ± 0.037 |
| | Female | 31.175 ± 3.65 | 0.478 ± 0.025 | 0.21 ± 0.051 | 0.237 ± 0.005 |
| 24 months | Male | 35.46 ± 2.33 | 0.446 ± 0.02 | 0.315 ± 0.035 | 0.245 ± 0.035 |
| | Female | 33.04 ± 3.16 | 0.47 ± 0.026 | 0.216 ± 0.032 | 0.266 ± 0.032 |

**Supplementary Table 1 | Weights of different organs of mice belonging to different age-groups.** The table contains the means and standard deviations of the body weights, weights of brain, heart and lungs of mice belonging to different age and sex. (n = 6 males and 6 females for the 2 months, 6 months, 12 months and 18 months-old cohorts ; n= 3 males and 3 females for the 24 months-old cohort).

| **Downregulated genes (46)** | **Upregulated genes (42)** |
|---|---|
| *Lims1, Tpst2, Htra1, Cnn3, Bag2, Pank1, Fyn, Leprotl1, Rhoj, Pdia6, Fermt2, Chek2, Tubb6, Kctd5, M6pr, Tm2d2, Tubb5, Tmem33, Xpnpep1, Sirt2, Dcaf12, Sh3gl1, Capzb, Slbp, Klf9, Nasp, Grn, Rell1, Ccdc85b, Tmem98, Gspt1, Slc31a1, Mrps7, Dapk3, Bcl10, Tpm1, Trerf1, Ddah2, Phc2, Map1lc3b, Cpne2, Mtmr14, Gnb1, Arf2, Cpt2, Rgl1* | *Zbtb20, Actg1, Fosl2, Ubr4, Trio, Mreg, Smg1, Iqgap1, Myo9a, Maml2, Zfc3h1, Cdc42bpa, Nbeal1, Rab11fip2, Med13l, Palld, Ell2, Ppp2r3a, Zmat1, Erc1, Nfkbiz, Junb, Tnrc6b, Spen, Aff1, Ppp3cb, Ep400, Sfi1, Ahnak, Ptgs2, Zswim6, Tmem117, Chd4, Birc6, Efemp2, Lpp, Myh9, Rev3l, Cdk14, Crim1, Stx11, Pla2g4a* |

**Supplementary Table 2 | SRF target genes significantly dysregulated with age (Linear regression analysis).** The table lists all the SRF target genes that were found to be significantly dysregulated with age in the cerebral ECs. Serum response factor (SRF) is a ubiquitous transcription factor that regulates the transcription of about 1000 genes. The list of ~ 1000 known SRF target genes identified in cultured NIH3T3 fibroblasts (Esnault et al., 2014) was used to identify the SRF target genes which were dysregulated with age in our RNA-seq dataset. The genes that were found to be significantly dysregulated with age (adjusted p-value $< 0.05$) in the linear regression analysis performed on all the samples, adjusted for sex, were considered. Out of 1388 genes that were significantly dysregulated with age, 88 genes (6.34 %) were SRF targets. Among them, 46 SRF target genes were found to be significantly downregulated, while 42 were significantly upregulated with age.

**Supplementary excel files.**

**Excel S1** List of all the differentially expressed genes in RNA-seq data (ordered according to increasing p-values).

**Excel S2** Geneset enrichment analysis listing all the differentially regulated pathways based on Top 1000 differentially expressed genes.

**Excel S3** Geneset enrichment analysis listing all the differentially regulated pathways based on Top 100 differentially expressed genes.

**Excel S4** Geneset enrichment analysis listing all the differentially regulated pathways based on Top 500 differentially expressed genes.

**Excel S5** List of genes/regions showing differential methylation status in cerebral ECs with age.

**Excel S6** List of regions and the genes annotated to the regions that show differential chromatin accessibility based on ATAC-seq data.

**Excel S7** List of common genes dysregulated with age in RNA-seq data and regions showing differential chromatin accessibility in ATAC-seq data.

References.

63. Weinl C, Riehle H, Park D, Stritt C, Beck S, Huber G, et al. Endothelial srf/mrtf ablation causes vascular disease phenotypes in murine retinae. *The Journal of clinical investigation*. 2013;123:2193-2206

64. Sivaraj KK, Dharmalingam B, Mohanakrishnan V, Jeong H-W, Kato K, Schröder S, et al. Yap1 and taz negatively control bone angiogenesis by limiting hypoxia-inducible factor signaling in endothelial cells. *Elife*. 2020;9:e50770

65. Buenrostro JD, Giresi PG, Zaba LC, Chang HY, Greenleaf WJ. Transposition of native chromatin for fast and sensitive epigenomic profiling of open chromatin, DNA-binding proteins and nucleosome position. *Nature methods*. 2013;10:1213-1218

66. Martin M. Cutadapt removes adapter sequences from high-throughput sequencing reads. *2011*. 2011;17:3

67. Dobin A, Davis CA, Schlesinger F, Drenkow J, Zaleski C, Jha S, et al. Star: Ultrafast universal rna-seq aligner. *Bioinformatics (Oxford, England)*. 2013;29:15-21

68. Ewels P, Magnusson M, Lundin S, Käller M. Multiqc: Summarize analysis results for multiple tools and samples in a single report. *Bioinformatics (Oxford, England)*. 2016;32:3047-3048

69. Love MI, Huber W, Anders S. Moderated estimation of fold change and dispersion for rna-seq data with deseq2. *Genome biology*. 2014;15:550

70. Marco-Sola S, Sammeth M, Guigo R, Ribeca P. The gem mapper: Fast, accurate and versatile alignment by filtration. *Nature methods*. 2012;9:1185-1188

71. Zhang Y, Liu T, Meyer CA, Eeckhoute J, Johnson DS, Bernstein BE, et al. Model-based analysis of chip-seq (macs). *Genome biology*. 2008;9:R137

72. Li H, Durbin R. Fast and accurate short read alignment with burrows-wheeler transform. *Bioinformatics (Oxford, England)*. 2009;25:1754-1760

73. Hovestadt V, Jones DT, Picelli S, Wang W, Kool M, Northcott PA, et al. Decoding the regulatory landscape of medulloblastoma using DNA methylation sequencing. *Nature*. 2014;510:537-541

74. Li H, Handsaker B, Wysoker A, Fennell T, Ruan J, Homer N, et al. The sequence alignment/map format and samtools. *Bioinformatics (Oxford, England)*. 2009;25:2078-2079

75. Liu Y, Siegmund KD, Laird PW, Berman BP. Bis-snp: Combined DNA methylation and snp calling for bisulfite-seq data. *Genome biology*. 2012;13:R61

76. Akalin A, Kormaksson M, Li S, Garrett-Bakelman FE, Figueroa ME, Melnick A, et al. Methylkit: A comprehensive r package for the analysis of genome-wide DNA methylation profiles. *Genome biology*. 2012;13:R87
